# Hoi1 targets the yeast BLTP2 protein to ER-PM contact sites to regulate lipid homeostasis

**DOI:** 10.1101/2025.02.11.637747

**Authors:** Samantha K. Dziurdzik, Vaishnavi Sridhar, Hailey Eng, Sarah D. Neuman, Junran Yan, Michael Davey, Stefan Taubert, Arash Bashirullah, Elizabeth Conibear

**Affiliations:** Centre for Molecular Medicine and Therapeutics, British Columbia Children’s Hospital Research Institute, University of British Columbia, Vancouver, Canada V5Z 4H4; Department of Medical Genetics, University of British Columbia, Vancouver, Canada V6H 3N1; Department of Cellular and Physiological Sciences, University of British Columbia, Vancouver, Canada V6H 3N1; Division of Pharmaceutical Sciences, University of Wisconsin-Madison, Madison, WI 53705-2222, USA

## Abstract

Membrane contact sites between organelles are important for maintaining cellular lipid homeostasis. Members of the recently identified family of bridge-like lipid transfer proteins (BLTPs) span opposing membranes at these contact sites to enable the rapid transfer of bulk lipids between organelles. While the VPS13 and ATG2 family members use organelle-specific adaptors for membrane targeting, the mechanisms that regulate other bridge-like transporters remain unknown. Here, we identify the conserved protein Ybl086c, which we name Hoi1 (Hob interactor 1), as an adaptor that targets the yeast BLTP2-like proteins Fmp27/Hob1 and Hob2 to ER-PM contact sites. Two separate Hoi1 domains interface with alpha-helical projections that decorate the central hydrophobic channel on Fmp27, and loss of these interactions disrupts cellular sterol homeostasis. The mutant phenotypes of BLTP2 and HOI1 orthologs indicate these proteins act in a shared pathway in worms and flies. Together, this suggests that Hoi1-mediated recruitment of BLTP2-like proteins represents an evolutionarily conserved mechanism for regulating lipid transport at membrane contact sites.

## Introduction

Eukaryotic cells maintain distinct organelle lipid compositions through both vesicular and non-vesicular transport pathways (Van Meer et al., 2008; Stefan et al., 2017). The non-vesicular pathway relies on lipid transfer proteins (LTPs) that move lipids between organelles at membrane contact sites (MCSs), regions where organelles are tethered together in close proximity (Prinz et al., 2020; Scorrano et al., 2019; Wong et al., 2019). Most LTPs use box-like domains to shuttle individual lipid molecules between membranes (Wong et al., 2019; Reinisch and Prinz, 2021). However, a recently identified class of elongated bridge-like lipid transfer proteins (BLTPs) spans opposing organelle membranes and forms a conduit for the bulk flow of lipids between them (Kumar et al., 2018; Maeda et al., 2019; Osawa et al., 2019; Valverde et al., 2019; Li et al., 2020, Neuman et al., 2021; Castro et al., 2022; Hanna et al. 2022; Toulmay et al., 2022). These BLTPs are thought to drive rapid membrane expansion, facilitate membrane repair, or promote large-scale changes in membrane composition in response to stress or other stimuli (reviewed in Neuman et al., 2022b and Hanna et al., 2023).

Vps13 and Atg2 were the first BLTPs to be structurally and biochemically characterized. Each has a hydrophobic groove that binds and transports glycerophospholipids (Kumar et al., 2018; Maeda et al., 2019; Osawa et al., 2019; Valverde et al., 2019; Li et al., 2020). Cryo-EM and structural prediction studies revealed that both proteins form rod-like structures, and the hydrophobic groove extends along the length of these structures to form a continuous lipid transfer channel (Chowdhury et al., 2018; Valverde et al., 2019; Maeda et al., 2019; Li et al., 2020). The groove consists of a repeating modular unit called the Repeating Beta Groove (RBG) domain, which contains several β-sheets followed by an overarching loop (Levine, 2022; Neuman et al., 2022b). The RBG domain defines a new superfamily of lipid transporters with long hydrophobic grooves that includes three additional, poorly-characterized members: BLTP1-3 (Braschi et al., 2022; Neuman et al., 2022b; Hanna et al., 2023).

To function as a bridge, each end of these proteins must target different membranes at a contact site. Vps13 and Atg2 rely on their N-terminus for ER targeting while organelle-specific adaptors recruit their C-terminus to acceptor membranes (Kotani et al., 2018; Maeda et al., 2019; Gómez-Sánchez et al., 2018; Park et al., 2013; John Peter et al., 2017; Bean et al., 2018; Guillén-Samander et al., 2021; Cai et al., 2022). ATG2 and the different VPS13 family members have characteristic domains, either inserted into the overarching loops or appended to the N or C-termini, that bind regulatory factors such as membrane-targeting adaptors and scramblases (Dziurdzik et al., 2020; Adlakha et al., 2022; Levine 2022). How other bridge-like lipid transporters are targeted to membrane contacts sites is unclear, as they often lack obvious regulatory domains and corresponding adaptors have not been identified.

The BLTP2-like lipid transport proteins are of particular interest, because they are conserved in eukaryotes yet their regulation and function is not well understood. The fly BLTP2 ortholog Hobbit (Hob) is present at ER-PM contacts and is required for regulated secretion events (Neuman and Bashirullah, 2018). Plant BLTP2 proteins are found at regions of the ER near the cell tip and cortex and are implicated in the growth of root tips, pollen tubes, and the cell plate during cell division (Aeschbacher et al., 1995; Procissi et al., 2003; Pietra et al., 2013; Pietra et al., 2015; Cheng and Bezanilla, 2021). The two BLTP2/Hobbit -like orthologs in yeast, Fmp27/Hob1 and Hob2, similarly localize to ER-PM contacts and may also be present at other contact sites (Neuman et al., 2022a; Toulmay et al., 2022), but how any BLTP2 ortholog is directed to these contacts is not known.

Here we investigate the role of the yeast Hob proteins, Fmp27/Hob1 and Hob2, at ER-PM contact sites. We identify Hoi1, a previously uncharacterized protein, as an adaptor that targets Fmp27 to the PM by binding alpha-helical projections extending from Fmp27’s central channel, an interaction that appears to be conserved in worms and flies. We show that the interaction between Hob proteins and Hoi1 is important for maintaining PM lipid homeostasis.

## Results

### Fmp27 shows saturable localization to ER-PM contact sites

AlphaFold predictions (Jumper et al., 2021) show that Fmp27 and Hob2 share a common architecture, with a series of β-sheets that form elongated hydrophobic channels characteristic of the RBG family, and a similar arrangement of extended α-helical protrusions (Fig. 1A). C-terminally tagged forms of Fmp27 and Hob2 localize to ER-PM contact sites, with Fmp27 being more readily detected due to its greater abundance (Neuman et al., 2022a; Toulmay et al., 2022). Tags at the C-terminus of Vps13 can selectively interfere with its distribution and function (Lang et al., 2015; Park et al., 2016). To further validate the endogenous localization of Fmp27, we used CRISPR/Cas9 to insert the fluorophore NeonGreen (NG) into a non-conserved loop after residue 737 (Fmp27^NG). Confocal imaging revealed Fmp27^NG at cortical sites that colocalized with an ER marker, Sec63-mScarletI, and showed partial overlap with the ER-PM contact site protein, Tcb3-mScarletI (Fig. 1B). A similar distribution was seen by widefield imaging when Fmp27 was tagged at the C-terminus with the bright GFP variant Envy (Fig. 1C), suggesting these tags do not perturb localization. This is consistent with observations that the C-terminally tagged form of the fly Fmp27 ortholog, Hobbit, is fully functional (Neuman et al., 2022a).

**Figure 1:**
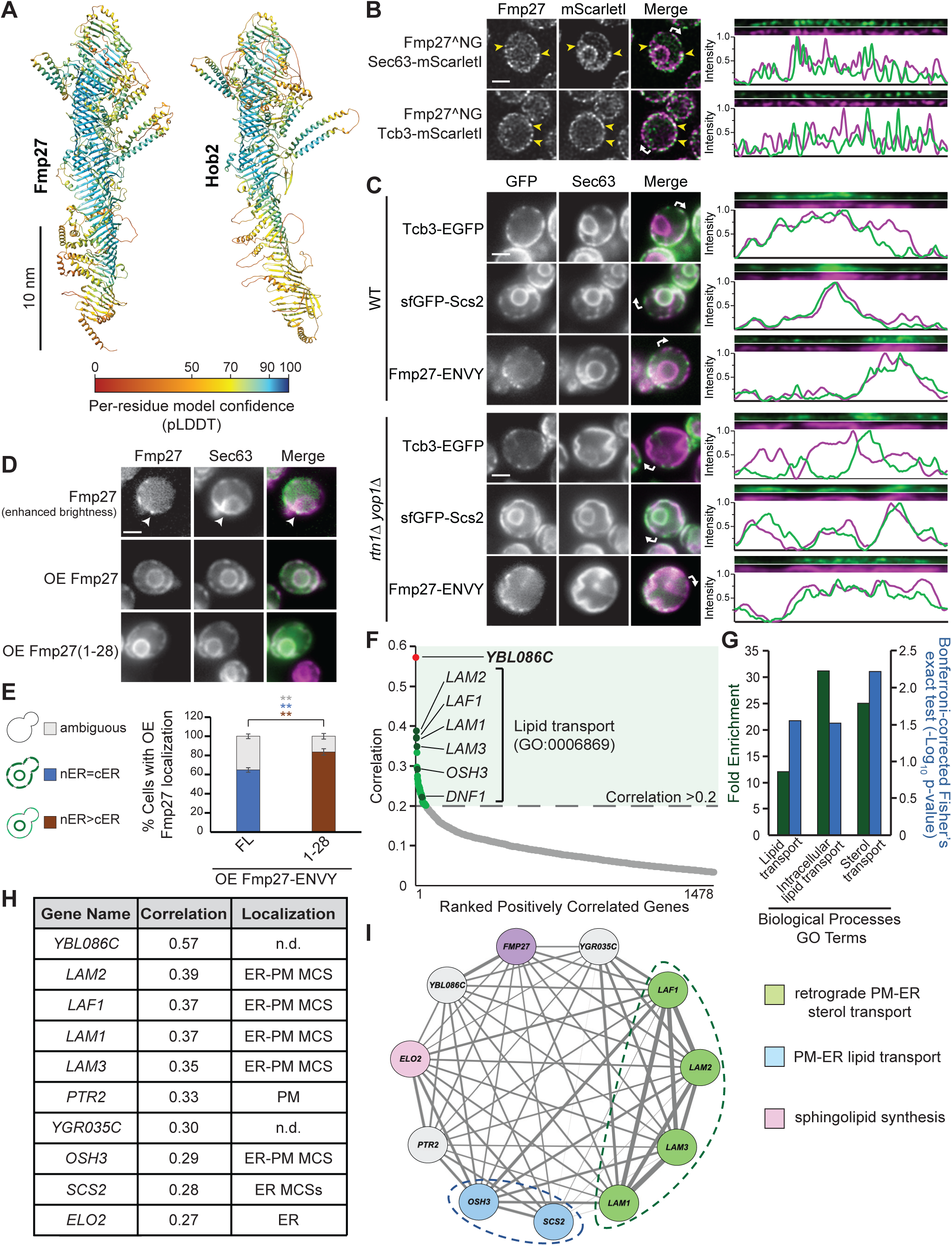
Fmp27 shows saturable localization to ER-PM contact sites. **(A)** Structural predictions of paralogous Fmp27 and Hob2 proteins acquired from the AlphaFold Protein Structure Database. Models are coloured by per-residue pLDDT values. Both proteins form elongated channels comprised of repeating ß-sheets and a similar arrangement of ⍺-helical protrusions. Bar = 10nm. **(B)** Confocal images of live yeast cells co-expressing Fmp27^NeonGreen (NG) and either the ER marker Sec63-mScarletI, or the ER-PM contact site marker TCB3-mScarletI, from their endogenous genomic loci. Linearized line scans of the cell cortex and normalized fluorescence signal intensities are shown to the right with curved arrows in the merged image indicating the line scan start site. Yellow arrowheads indicate sites of colocalization. Bar = 2µm. **(C)** Fmp27-ENVY maintains colocalization with Sec63-mScarletI in *rtn1*Δ *yop1*Δ cells. Widefield images of wild-type and *rtn1*Δ *yop1*Δ yeast cells expressing Sec63-mScarletI and either Tcb3-EGFP, sfGFP-Scs2 or Fmp27-ENVY, from their endogenous genomic loci. Linearized line scans of the cell cortex and normalized fluorescence intensities are shown to the right with curved arrows in the merged image indicating the line scan start site. Bar = 2µm. (**D**) Localization of Fmp27 at cortical patches is saturable. On overexpression, Fmp27-Envy localizes throughout the ER, marked by Sec63-mScarletI. Fmp27 residues 1-28 are sufficient for ER localization. The image of Fmp27-Envy expressed at endogenous levels is shown at enhanced brightness so the localization can be seen. White arrowheads indicate Fmp27-ENVY localization at ER-PM contacts. Scale bar =2µm. (**E**) Quantification of the ER localization of the overexpressed full length Fmp27-Envy (FL), or the overexpressed N-terminal fragment Fmp27^1-28^-Envy by blind scoring of cells in D. Unpaired two-tailed t-test with Welch’s correction; >100 cells/replicate/strain, n=3, p**<0.01. Color of asterisks corresponds to the significance of each localization category. Error bars indicate SEM. nER=nuclear ER; cER = cortical ER. **(F)** Ranked genes positively correlated with *FMP27* in the homozygous deletion profiling (HOP) chemogenomic dataset (Hoepfner et al., 2014) with p-values <0.01. Genes with correlation scores >0.2 identified by the Lipid Transport GO term (GO:0032365) are labeled (dark green dots). *YBL086C* (red dot) is the gene with the highest positive correlation to *FMP27*. **(G)** Biological process Gene Ontology (GO) functional enrichment analysis for genes from F with correlations >0.2 (http://geneontology.org). GO terms are presented with their fold enrichment (green) and negative log_10_ p-values from Bonferroni-corrected Fischer’s exact test (blue). Sterol Transport (GO:0015918), Lipid Transport (GO:0006869) and Intracellular Lipid Transport (GO:0032365) are significantly enriched ontologies. **(H)** Table of the 10 most highly correlated genes from F with their respective *FMP27* correlation values and observed localizations listed. **(I)** Network of *FMP27* and highly correlated genes from H. Edges indicate the correlation score between nodes with thicker edges indicating higher correlation values. Nodes are colored according to known roles in lipid transport. Genes with unknown functions are shown in grey, with *FMP27* in purple. Functionally related genes are denoted with dashed lines.

The partial colocalization of Fmp27 with Tcb3 suggests that these proteins may not entirely reside at the same ER-PM contacts. Tcb3 localizes to sites of PM contact with tubular ER, whereas the VAP homolog Scs2 is preferentially found at sites of PM contact with flat ER sheets (Hoffmann et al., 2019; Collado et al., 2019). These classes of ER-PM contact sites can be distinguished in cells lacking the ER-tubulating reticulon proteins Rtn1 and Yop1 (Hoffmann et al., 2019). We found that Fmp27-ENVY, like sfGFP-Scs2, colocalizes with Sec63-mScarletI at the expanded ER sheets found in *rtn1Δ yop1Δ* mutants, while Tcb3-EGFP remains at cortical sites, which represent contacts with residual tubular ER (Fig. 1C; Hoffmann et al., 2019). These results suggest that Fmp27 and Tcb3 predominantly occupy distinct populations of ER-PM contacts.

When overexpressed, Fmp27-ENVY redistributed from cortical sites and was found along the entire ER network (Fig. 1D). Since the Fmp27 N-terminus is necessary for ER localization and protein stability (Neuman et al., 2022a), we tested if it is sufficient for ER targeting. Indeed, the first 28 residues of Fmp27 fused to GFP localized to the ER (Fig. 1D-E), suggesting that Fmp27 is tethered to the ER by its N-terminal helix (Fig. 1A). We speculated that Fmp27 localization at ER-PM contacts requires binding of the Fmp27 C-terminus to a partner at the PM, and that this interaction is saturated when Fmp27 is overexpressed, resulting in its redistribution along the ER.

To uncover candidate PM targeting factors for Fmp27, we used a chemogenomic profiling approach to identify genes that are functionally linked to *FMP27* (Fig. 1F-I). Pearson correlation coefficients were computed from a large-scale dataset that contains the sensitivity scores of all non-essential homozygous deletion mutants exposed to 1800 compounds (Hoepfner et al., 2014). Gene Ontology (GO) functional enrichment analysis (geneontology.org) of genes that showed a positive correlation with *FMP27* identified Sterol Transport (GO:0015918), Lipid Transport (GO:0006869) and Intracellular Lipid Transport (GO:0032365) as significantly enriched ontologies, supporting a role for Fmp27 in lipid transport (Fig. 1F,G). The top ten correlated genes included a number of ER-PM contact site proteins and an ER-localized lipid biosynthetic enzyme (Fig. 1H). Network analysis of *FMP27* and these 10 genes revealed that three Lam family proteins, which are involved in retrograde PM-to-ER transport, form a distinct cluster together with the Lam2/4-associated regulatory factor Laf1 (Gatta et al., 2015; Murley et al., 2015; Topolska et al., 2020; Fig. 1I). Notably, the uncharacterized gene *YBL086C* showed the strongest reciprocal correlation with *FMP27*. We named this gene *HOI1* (for HOb Interactor 1) and selected it for further characterization.

### The uncharacterized protein Hoi1 recruits Fmp27 to the PM

Hoi1 has a predicted N-terminal C2 (NT-C2) domain that could mediate lipid binding at membranes (Zhang and Aravind, 2010) and a long relatively unstructured C-terminus (Fig. 2A). To determine if Hoi1 recruits Fmp27 to the PM, we tested if increased *HOI1* expression could restore overexpressed Fmp27^NG to ER-PM contact sites (Fig. 2B-C). Indeed, when both Hoi1 and Fmp27 were expressed from the *ADH1* promoter, Fmp27^NG was no longer distributed along the nuclear and cortical ER, but instead showed enhanced localization to cortical sites at the PM. Consistent with a targeting role for Hoi1, epitope-tagged versions of Fmp27 and Hoi1 showed a striking co-localization at cortical patches when co-overexpressed (Fig. 2D).

**Figure 2.**
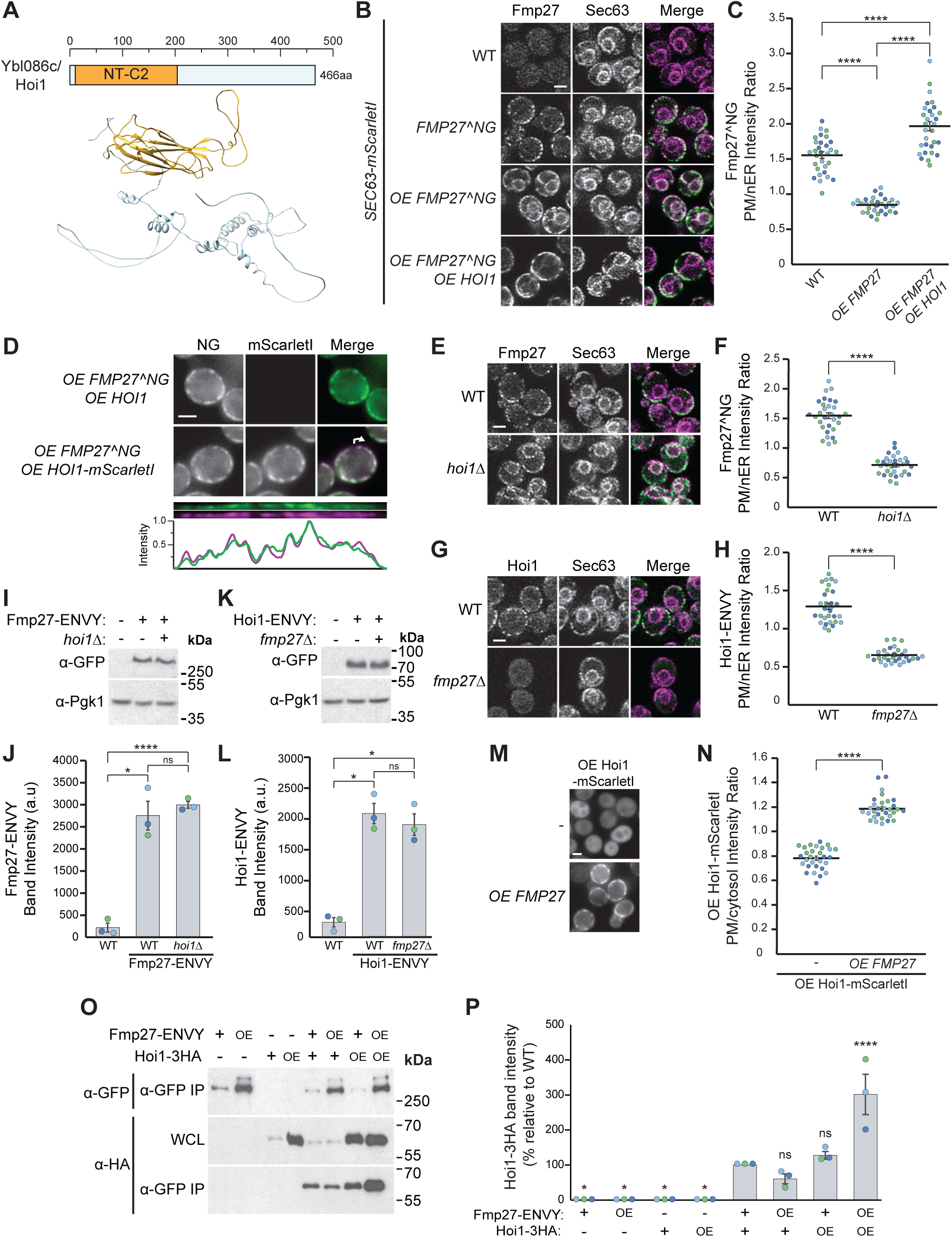
The uncharacterized protein Hoi1 is an Fmp27 binding partner at the PM. **(A)** Domain schematic and predicted structure of Ybl086c/Hoi1 from the AlphaFold Protein Structure Database (https://alphafold.ebi.ac.uk/; Jumper et al., 2021). The annotated NT-C2 domain is colored gold. Scale shows number of amino acids. **(B)** Overexpression (OE) of *HOI1* restores the ER-PM localization of overexpressed Fmp27^NeonGreen (NG). Confocal images of yeast cells expressing Fmp27^NG and Sec63-mScarletI from their genomic loci either from their native promoter or the strong *ADH1* promoter. Bar = 2µm. **(C)** Quantitation of the PM to nuclear ER intensity ratio of Fmp27^NeonGreen (NG) from yeast strains in B determined with line scans. The average intensity ratio is presented. Individual data points are overlaid and colored by replicate. One-way ANOVA with Tukey’s multiple comparison test; n = 3; cells/replicate/strain = 10; **** = p<0.0001. Error bars indicate SEM. **(D)** Overexpressed (OE) Fmp27^NeonGreen (NG) colocalizes with overexpressed Hoi1-mScarletI at cortical sites. Widefield images of yeast with a linearized line scan and normalized fluorescent signal intensities shown below. The curved arrow in the merged image indicates the line scan start site. Bar = 2µm. **(E)** Fmp27^NeonGreen relies on *HOI1* for recruitment to the PM. Confocal images of wild-type or *hoi1*Δ yeast cells expressing Fmp27^NeonGreen and Sec63-mScarletI from their endogenous genomic loci. Bar = 2µm. **(F)** Quantitation of the PM to nuclear ER intensity ratio of Fmp27^NeonGreen (NG) from yeast strains in E determined from line scans. The average intensity ratio is presented. Individual data points are overlaid and colored by replicate. Non-parametric two-way t-test; n = 3, cells/replicate/strain = 10, **** = p<0.0001. Error bars indicate SEM. **(G)** Hoi1-ENVY relies on *FMP27* for localization to the PM. Confocal images of wild-type or *fmp27*Δ yeast cells expressing Hoi1-ENVY and Sec63-mScarletI from their endogenous genomic loci. Bar = 2µm. **(H)** Quantitation of the PM to nuclear ER intensity ratio of Hoi1-ENVY from yeast strains in G determined with line scans. The average intensity ratio is presented. Individual data points are overlaid and colored by replicate. Non-parametric two-way t-test; n = 3, cells/replicate/strain = 10, **** = p<0.0001. Error bars indicate SEM. **(I)** Fmp27-ENVY stability is unaffected by deletion of *HOI1*. Anti-GFP *w*estern blot of Fmp27-ENVY in wild-type and *hoi1*Δ yeast strains. Pgk1 serves as a loading control. **(J)** Quantitation of Fmp27-ENVY band intensities in I by densitometry. Individual data points are overlaid and colored by replicate. One-way ANOVA with Tukey’s multiple comparison test; n = 3, **** = p<0.0001, * = p<0.5, ns = not significant. Error bars indicate SEM. **(K)** Hoi1-ENVY stability is unaffected by deletion of *FMP27*. Anti-GFP *w*estern blot of Hoi1-ENVY in wild-type and *fmp27Δ* yeast strains. Pgk1 serves as a loading control. **(L)** Quantitation of Hoi1-ENVY band intensities in K by densitometry. Individual data points are overlaid and colored by replicate. One-way ANOVA with Tukey’s multiple comparison test; n = 3, * = p<0.5, ns = not significant. Error bars indicate SEM. **(M)** Overexpressed (OE) *FMP27* restores PM localization of overexpressed Hoi1-mScarletI. Widefield images of yeast cells overexpressing Hoi1-mScarletI with endogenous or overexpressed *FMP27*. Bar = 2µm. **(N)** Quantitation of the PM to cytosol intensity ratio of Hoi1-mScarletI from yeast strains in M determined from line scans. The average intensity ratio is presented. Individual data points are overlaid and colored by replicate. Non-parametric two-way t-test; n = 3, cells/replicate/strain = 10, **** = p<0.0001. Error bars indicate SEM. **(O)** Immunoprecipitation (IP) of ENVY-tagged Fmp27 co-purifies 3HA-tagged Hoi1. Tagged proteins are expressed from their endogenous genomic loci, either at native levels or overexpressed (OE) from a strong *ADH1* promoter. WCL: whole cell lysate. **(P)** Quantification of the interaction between Fmp27-ENVY and Hoi1-3HA in strains shown in O by densitometry. One-way ANOVA with Dunnett’s correction comparing the interaction between Fmp27-ENVY and Hoi1-3HA expressed at native levels to other strains; n=3, p*<0.5, p****<0.0001. Error bars indicate SEM. Only significant differences are indicated. OE: overexpressed.

In cells lacking Hoi1, endogenous Fmp27^NG was no longer enriched at cortical puncta but instead redistributed along the entire ER (Fig. 2E-F), indicating that Hoi1 is required for Fmp27 localization to the PM. We found that endogenously expressed Hoi1-ENVY also localized to cortical puncta (Fig. 2G). Unexpectedly, in *fmp27Δ* mutants Hoi1-ENVY lost its PM localization and became cytosolic (Fig. 2G-H). This change in Hoi1 localization was not due to altered protein levels, as Western blotting showed that neither protein’s abundance depends on the presence of the other (Fig. I-L). These results reveal an unexpected reciprocal requirement, where Fmp27 is needed for Hoi1’s PM localization. Indeed, Hoi1-mScarletI was largely cytosolic when overexpressed, although some dim cortical puncta could also be observed, and high levels of Fmp27 restored the PM localization of overexpressed Hoi1-mScarletI (Fig. 2M-N). Together, this indicates that the PM recruitment of both proteins is interdependent and suggests that Fmp27 and Hoi1 might target the PM as a complex.

To test if Fmp27 and Hoi1 physically interact, we performed co-immunoprecipitations (coIPs) with endogenous or overexpressed tagged forms of each protein. Hoi1-HA copurified with Fmp27-ENVY and the recovery of both proteins increased only when both proteins were overexpressed (Fig. 2O-P). This enhanced recovery suggests either that the proteins directly interact, or that the abundance of any bridging protein is not limiting for complex formation.

Collectively, these results identify Hoi1 as a Fmp27 interactor and show that both proteins rely on each other for recruitment to ER-PM contact sites.

### The C-terminus of Hoi1 recruits Fmp27 to PM contact sites

We used co-IP to identify the region of Hoi1 that interacts with Fmp27. Fmp27 co-purified with the unstructured C-terminus of Hoi1 (Hoi1^216-466^-ENVY), but not with the NT-C2 domain-containing fragment (Hoi1^1-216^-ENVY) (Fig. 3A-B). The Fmp27 paralog Hob2 showed the same binding pattern, co-purifying with both full length Hoi1and its C-terminal domain (Fig. 3C-D). These results indicate that Hoi1 interacts with both Fmp27 and Hob2 via its C-terminus.

**Figure 3:**
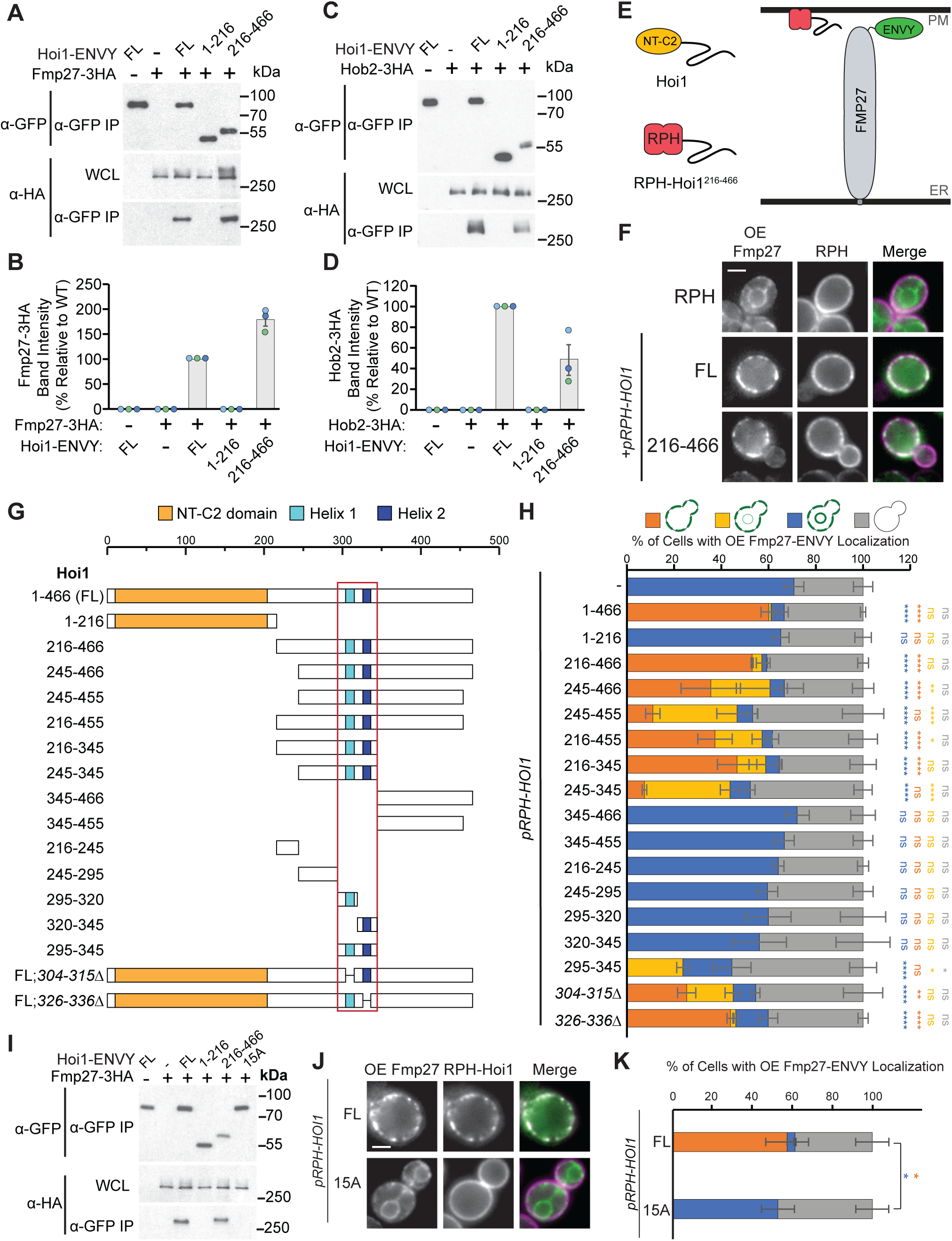
The C-terminus of Hoi1 recruits Fmp27 to PM contact sites. **(A)** The Hoi1 tail is necessary and sufficient for Fmp27 binding. Anti-GFP IP of plasmid-expressed full-length (FL) Hoi1-ENVY, Hoi1 N-terminus (1-216)-ENVY or C-terminus (216-466)-ENVY from strains co-expressing Fmp27-3HA from the endogenous locus. WCL: whole cell lysate. (**B**) Quantification of co-purifying Fmp27-3HA by densitometry of samples shown in A. The % band intensity is relative to the amount of Fmp27-3HA that coprecipitated with full length (FL) Hoi1-ENVY, n=3. **(C)** The Hoi1 tail, but not the Hoi1 N-terminal domain, also binds Hob2. Anti-GFP IP of plasmid-expressed full-length (FL) Hoi1-ENVY, Hoi1 N-terminus (1-216)-ENVY, or C-terminus (216-466)-ENVY from strains co-expressing genomically tagged Hob2-3HA. WCL: whole cell lysate. **(D)** Quantification of co-purifying Hob2-3HA by densitometry of samples shown in C. The % band intensity is relative to the amount of Hob2-3HA coprecipitated with full-length (FL) Hoi1-ENVY, n=3. **(E)** Schematic of the Fmp27 recruitment assay. The putative membrane-binding NT-C2 domain of Hoi1 is replaced by the fluorescently labelled PM marker, Cherry-2xPH^PLCδ^ (RPH), which enhances PM targeting. (**F**) Overexpressed Fmp27-ENVY is recruited to cortical sites by full length Hoi1, or the C-terminal tail (216-466) of Hoi1, fused to the RPH module. Scale bar =2µm. **(G)** Schematic of Hoi1 fragments tested for Fmp27 recruitment. All Hoi1 fragments were expressed from plasmids as RPH chimeras. The red box indicates the minimal region of Hoi1 that confers Fmp27 recruitment. (**H**) The PM recruitment of overexpressed Fmp27-ENVY by different RPH-Hoi1 chimeras relative to RPH alone. Images were blinded and scored for relative intensity of PM to ER labeling. Dark orange: PM only; light orange: PM>ER; blue: PM=ER; grey: no clear localization. One-way ANOVA with Dunnett’s correction; n=3, cells/strain/replicate>100, p*<0.05, p**<0.01, p***<0.001, p****<0.0001. Error bars indicate SEM. Only significant differences are indicated. **(I)** Anti-GFP IP of plasmid-expressed, ENVY-tagged forms of full-length (FL) Hoi1-ENVY, Hoi1 N-terminus (1-216), Hoi1 C-terminus (216-466) or the Hoi1-15A mutant from strains co-expressing genomically tagged Fmp27-3HA. WCL: whole cell lysate. (**J**) Overexpressed Fmp27-ENVY is recruited to cortical sites by the full length (FL) Hoi1 fused to RPH, but not by a mutant version where 15 conserved residues in the Hoi1 C-terminus were mutated to alanine (15A). Scale bar =2µm. (**K**) Quantification of strains shown in J was carried out as described above in H. One-way ANOVA with Dunnett’s correction; n=3, cells/strain/replicate>100, p*<0.05. Error bars indicate SEM.

Although the Hoi1 C-terminus binds Fmp27, this fragment did not localize to cortical ER-PM sites when co-overexpressed with Fmp27. Instead, it distributed evenly along the ER in a Fmp27-dependent manner (Fig. S1A-B). This suggests the NT-C2 domain is important for PM targeting. However, the Hoi1 NT-C2 domain, when expressed on its own, was unable to localize to membranes in either WT or *FMP27*-overexpressing cells (Fig. S1A-B). Thus, while the Hoi1 C-terminus is necessary and sufficient for binding both Fmp27 and Hob2, proper targeting to cortical sites requires the complete protein.

To better understand domain requirements for PM targeting, we created chimeric proteins where the poorly targeted NT-C2 domain of Hoi1 was replaced with a PM-localized “RPH” reporter module consisting of the <u>R</u>FP variant mCherry fused to the tandem <u>PH</u> domains of PLCδ, which bind PM-enriched pools of PI(4,5)P_2_ (Stauffer et al., 1998; Fig. 3E). When either Hoi1 or its C-terminal tail were fused to the RPH module, the resulting chimeric proteins strongly recruited overexpressed Fmp27-ENVY (Fig. 3F) or Hob2-Envy (Fig. S1C) to cortical PM patches. We tested whether the NT-C2 domain could be co-recruited to these Fmp27/RPH-Hoi1 assemblies, but found it remained cytosolic even when Fmp27 was tethered to the PM by the RPH-Hoi1^216-466^ chimera (Fig. S1D,E). This suggests that the Hoi1 NT-C2 domain must be linked to its C-terminal tail for PM targeting.

To narrow down the Fmp27 binding site on Hoi1, we generated a series of RPH chimeras containing different regions of the Hoi1 C-terminus and tested their ability to recruit overexpressed Fmp27-ENVY to cortical sites (Fig. 3G-H). The smallest fragment retaining recruitment activity corresponded to residues 295-345, which contains two highly conserved helices: Helix 1 (residues 304-315) and Helix 2 (residues 326-336) (Fig. S1F-G). Since neither helix alone was necessary or sufficient for Fmp27 recruitment, we generated a mutant form of Hoi1 where 15 conserved residues within or flanking these helices were replaced with alanines (Fig. S1G). This Hoi1^15A^ mutant was stably expressed, but did not coIP with Fmp27 (Fig. 3I; Fig. S1H), nor recruit overexpressed Fmp27-ENVY to cortical sites when fused to the RPH module (Fig. 3J-K). Based on these results, we conclude that conserved residues within and adjacent to the two helices are necessary for Fmp27 interaction and membrane recruitment.

### Identifying the interacting regions of Fmp27 and Hoi1

To find the region of Fmp27 that interacts with Hoi1, we generated a series of Fmp27-ENVY C-terminal truncations (Fig. 4A) and tested their recruitment to cortical sites by the RPH-Hoi1 chimera (Fig. 4B-D). Many Fmp27 truncation mutants, which lacked sections of the hydrophobic groove, formed bright aggregates when overexpressed (Fig. 4B). Strikingly, co-overexpression of the RPH-Hoi1 chimera blocked aggregation and restored some degree of cortical PM localization for truncations containing at least the first 1834 residues of Fmp27 (Fig. 4B-D). Shorter constructs formed aggregates that were not recruited by RPH-Hoi1, which could reflect either loss of interacting residues or large-scale misfolding that impacts a distant binding interface. These results indicate the C-terminal 794 residues of Fmp27 are largely dispensable for recruitment by the Hoi1 chimera (Fig. 4E), and identify the region of Fmp27 that is minimally sufficient for the Hoi1 interaction.

**Figure 4.**
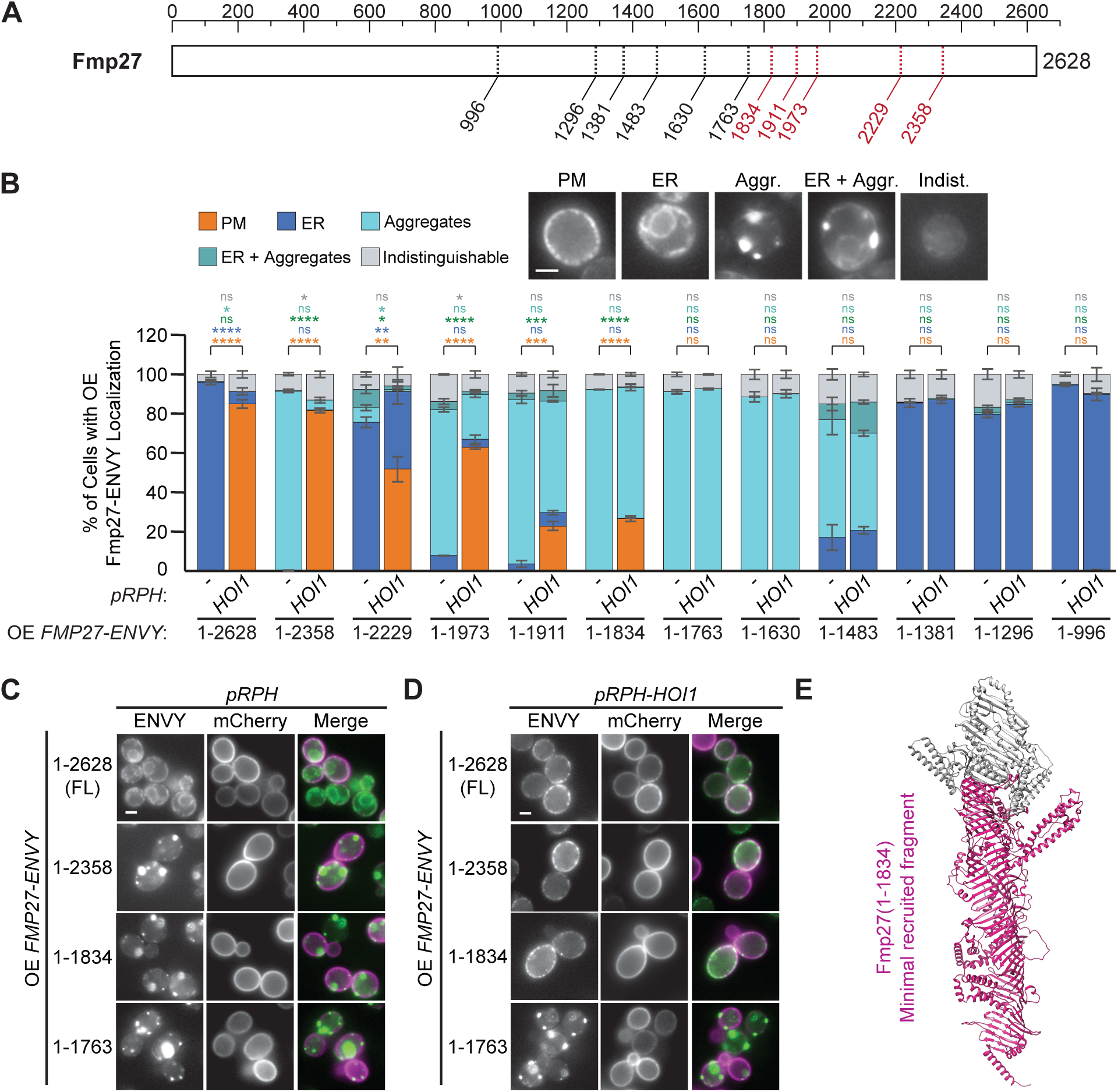
Recruitment by Hoi1 does not require the Fmp27 C-terminus. **(A)** Schematic of Fmp27 indicating the C-terminal truncations tested for PM recruitment by the plasmid-expressed RPH-Hoi1 chimera. Truncations that were identified as sufficient for recruitment are labelled in red. **(B)** Quantitation of the localization of overexpressed (OE) Fmp27-ENVY truncations in live yeast cells expressing the RPH module alone, or the RPH-Hoi1 chimera, by counting of blinded images. Representative images for each localization category are shown. Bar = 2µm. Unpaired two-tailed parametric T-tests; n=3, cells/strain/replicate ≥ 306; **** = p<0.0001, *** = p<0.001, ** = p<0.01, * = p<0.5, ns = not significant. Error bars indicate SEM. **(C, D)** Representative widefield images from select strains in B of overexpressed (OE) Fmp27-ENVY truncations in cells expressing (C) RPH alone, or (D) the RPH-Hoi1 chimera. Bar = 2µm. **(E)** AlphaFold predicted structure of Fmp27 (Jumper et al., 2021) indicating the minimal fragment that confers Hoi1 recruitment in magenta.

To identify the Fmp27-Hoi1 binding interface we modeled interactions between full-length Hoi1 and overlapping fragments of Fmp27 using the AlphaFold2-based ColabFold algorithm (Jumper et al., 2021; Mirdita et al., 2022). We removed low-scoring regions that typically represent disordered regions, considering only modeled residues with a predicted local distance difference test (pLDDT) score>70, which indicates a generally correct backbone placement (Tunyasuvunakool et al., 2021). This identified a confident interface between Hoi1 and the C-terminal third of Fmp27 (Fig. 5A-B, Fig. S2A-B), which buries a surface of 2622.6 Å^2^ (for PISA analysis see Table S1).

**Figure 5:**
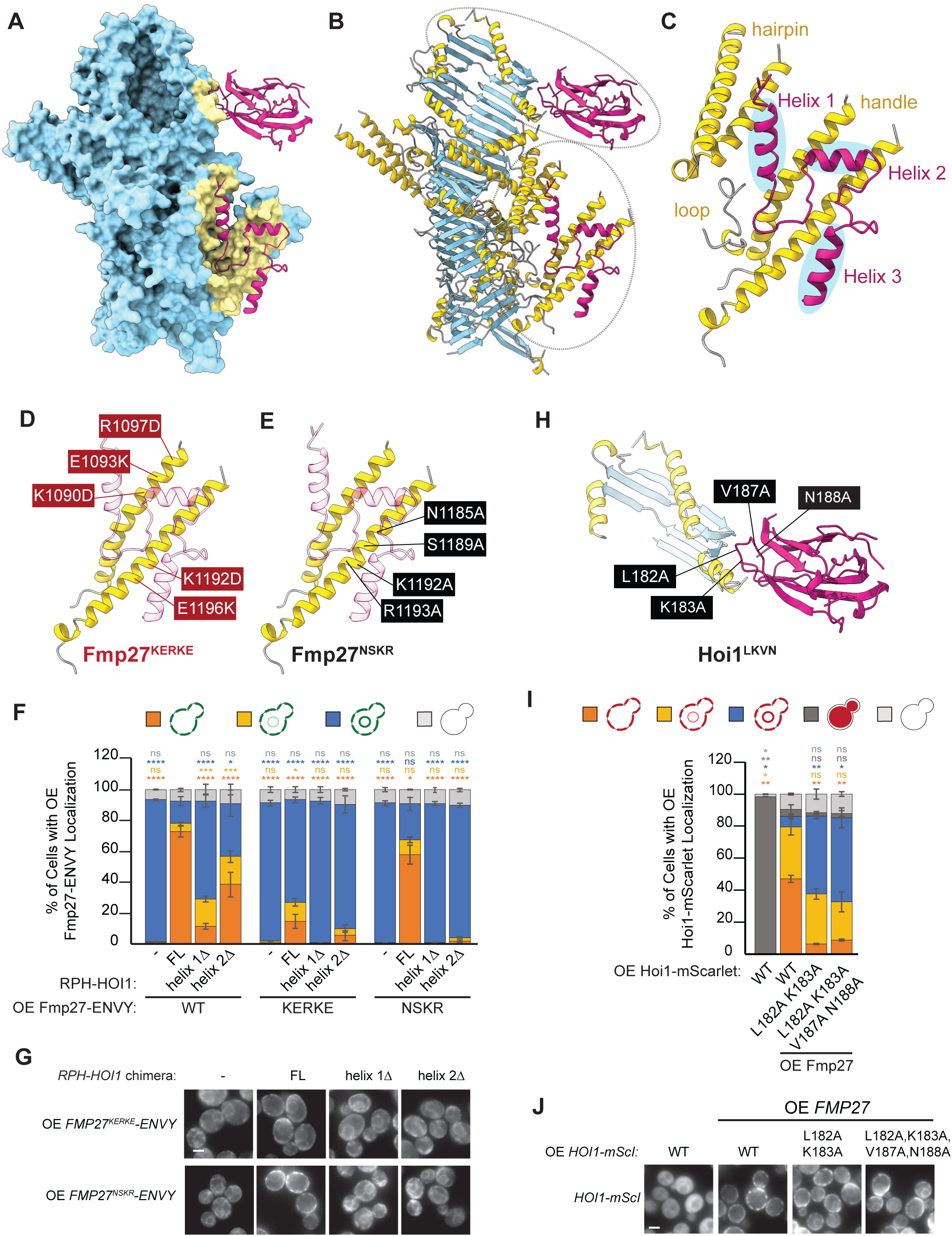
AlphaFold identifies a predicted Fmp27/Hoi1 interface. **(A,B)** ColabFold-predicted interaction between the Fmp27 C-terminus (residues 1051-2628) and Hoi1. The top-scoring model, showing residues with pLDDT values >70, is illustrated. (A) shows a surface view of Fmp27 (blue) with Hoi1 in magenta, and the Fmp27 interface residues in yellow. (B) shows ribbon views of both partners. Fmp27 α-helices: yellow; Fmp27 β-sheets: blue; Hoi1: magenta. The circles indicate the regions shown in panels C and H. **(C)** Close up of interacting regions of Fmp27 (yellow) and the Hoi1 C-terminal tail (magenta). The region of Hoi1 that includes Helices 1-3 and their flanking sequences interacts with three structural elements in Fmp27: the “handle” which consists of two interacting helices (residues 1065-1216), an unstructured “loop” (residues 1524-1540), and a helical ‘hairpin” (residues 2272-2330). **(D)** Schematic indicating the Fmp27 interface residues substituted for lysine or aspartic acid in the Fmp27^KERKE^ mutant. **(E)** Schematic indicating the Fmp27 interface residues substituted for alanine in the Fmp27^NSKR^ mutant. **(F)** Mutating Fmp27 interface residues disrupts Hoi1-mediated recruitment. The localization of overexpressed (OE) WT, Fmp27^KERKE^ or Fmp27^NSKR^ mutant forms of Fmp27-ENVY, in cells co-overexpressing RPH-Hoi1 chimeras, was quantified by counting of blinded images. Cells/strain/replicate ≥ 215; n = 3. One-way ANOVA with Dunnett’s corrected post-hoc test comparing to WT Fmp27 recruited by the WT RPH-Hoi1 chimera; **** = p<0.0001; *** = p<0.001; * = p<0.5; ns = not significant. Error bars indicate SEM. The Hoi1 residues removed in the *helix 1*Δ chimera are 304-315 and the residues removed in *helix 2*Δ are 326-336. **(G)** Representative widefield images of strains quantified in F. Bar = 2µm. **(H)** Close up of interacting regions of Fmp27 (helices, yellow; sheets, blue) and the Hoi1 N-terminal C2 domain (magenta). Hoi1 interface residues substituted for alanine in the Hoi1^LKVN^ mutant are indicated. **(I)** Mutation of Hoi1 NT-C2 interface residues causes the *FMP27*-dependent re-localization of Hoi1 to the ER. Wild type (WT) or mutant forms of mScarlet-tagged Hoi1 were overexpressed in cells that were co-overexpressing untagged *FMP27*. One-way ANOVA with Dunnett’s multiple comparisons test; n=3, cells/strain/replicate >130, ** = p<0.01, * = p<0.5, ns = not significant. Error bars indicate SEM. **(J)** Representative widefield images of the strains quantified in I. Bar = 2µm.

In this model, Fmp27 contacts residues 297-362 of the Hoi1 tail (Fig. 5C). This region encompasses Helix 1 and 2 with their flanking sequences, which are part of the minimal recruitment region (see Fig. 3G, Fig. S1G), and a C-terminal “Helix 3” that was largely dispensable in our recruitment assay. Several metrics support the validity of this predicted interface (Fig. S2): the predicted aligned error (PAE) values between interface residues are low (<5 Å), the pLDDT scores are high (>70), and importantly, 13 of the 15 conserved Hoi1 residues that we showed were essential for Fmp27 binding (Fig. 3I, Fig. S1F-H) lie within the predicted interface (Table S1).

The AlphaFold2 model revealed that the Hoi1 C-terminal region, including Helices 1-3 and their surrounding residues, contacts a prominent pair of long intertwined α-helices that project from the central hydrophobic channel of Fmp27, which we refer to as the “handle” (Fig. 4E, Fig. 5C). The Hoi1 critical region also contacts a second pair of Fmp27 helices that are linked by a hairpin turn, and an intervening loop. This structural prediction aligns with our truncation analysis (Fig. 4B), where removing the hairpin helices (Fmp27^1-2229^) partially reduced Hoi1-mediated recruitment. Complete loss of recruitment occurred upon deletion of Fmp27 residues 1803-1807, which are predicted to stabilize the intervening loop and handle.

To test if the handle is necessary for Hoi1 binding, we initially used CRISPR/Cas9 mutagenesis to either replace it entirely, or mutate multiple Hoi1-interacting residues. These Fmp27 mutants aggregated on overexpression and were not recruited to the PM by the RPH-Hoi1 chimera. By mutating subsets of the predicted interacting residues, we generated two Fmp27 mutants that did not aggregate and, like wild-type Fmp27, localized to the ER on overexpression. The first mutant, Fmp27^KERKE^, has 5 charge-swapped substitutions that are predicted to disrupt a major portion of the handle’s interface with Hoi1 (Fig. 5D). This mutant showed severely impaired PM recruitment by RPH-Hoi1 (Fig. 5F,G), confirming the handle’s essential role in Hoi1-mediated targeting.

The model also predicts contributions from residues connecting the Hoi1 helices. To probe these interactions, we examined a second mutant, Fmp27^NSKR^, that has alanine substitutions in 4 residues that are predicted to contact regions flanking Hoi1’s helix 2 and at helix 3, but not helix 2 itself (Fig. 5E). While this mutant showed normal recruitment by wild-type RPH-Hoi1, it failed to respond to an RPH-Hoi1 variant lacking helix 2 (Fig. 5F,G). This synergistic effect suggests that Hoi1 helix 2 and its flanking regions cooperate to create a stable interface with the Fmp27 handle. Together, these experiments validate the AlphaFold2-predicted interface and establish it as the primary binding site for the Hoi1 C-terminal tail.

Unexpectedly, the AlphaFold2 model also predicted a confident interface between Fmp27 and conserved loops in the Hoi1 NT-C2 domain (Fig. 5A,B,H). This interface is considerably smaller than that involving the Hoi1 C-terminal tail (620.9 Å vs 2001.7 Å), and our coIP and fluorescence imaging experiments showed the NT-C2 domain is not sufficient to drive Fmp27 binding (Fig. 3A) or membrane recruitment (Fig. 3G-H). However, mutating residues in a conserved NT-C2 loop that is predicted to contact a helix in Fmp27 significantly impaired the Fmp27-dependent localization of Hoi1 to the plasma membrane (Fig. 5I,J), suggesting this interface helps stabilize the PM targeting of the Fmp27-Hoi1 complex.

Our comprehensive mutational analysis demonstrates that Hoi1-mediated PM recruitment primarily depends on an Fmp27’s prominent α-helical extension, with additional contributions from nearby loop and helical regions. A direct interaction between the Hoi1 NT-C2 domain and the Fmp27 C-terminus also contributes to the stable association of the complex with the plasma membrane.

### Fmp27 and Hoi1 have a shared role in lipid homeostasis

What is the function of the Fmp27-Hoi1 complex at ER-PM contacts? Our analysis of correlated chemogenomic profiles suggested both proteins are functionally linked to members of the LAM/GRAMD family of sterol transporters at these contact sites (Fig. 1I). *LAF1* is required for the LAM-mediated retrograde transport of ergosterol from the PM to the ER (Topolska et al., 2020), and knockout of this gene causes ergosterol accumulation at the PM and sensitivity to Amphotericin B (AmB), which binds accessible ergosterol on the outer PM leaflet (Parsons et al., 2006; Gray et al., 2012). Large-scale drug sensitivity screens also identified *fmp27* and *hoi1* mutants as AmB sensitive (Parsons et al., 2006), supporting a shared role in maintaining ergosterol homeostasis.

We used a plate reader assay to quantify the drug sensitivity of mutant strains grown in rich yeast peptone dextrose (YPD) media with a sublethal concentration of AmB (0.4 μM) (Fig. 6A-C). While the AmB sensitivity of the *fmp27Δ* single mutant was not significantly different from the wild type strain, the double *fmp27*Δ *hob2*Δ mutant displayed a sensitivity that was similar to that of a *hoi1*Δ or *hoi1*Δ *fmp27*Δ mutant. These results, confirmed by spot assays (Fig. S3A), suggest that Fmp27 and its paralog Hob2 have redundant functions and act together with Hoi1. We used this AmB sensitivity phenotype to show that the internally tagged Fmp27^NG was fully functional (Fig. S3B-D) and to test the functional importance of the Fmp27-Hoi1interaction (Fig. 6D-F). Both untagged and ENVY-tagged *HOI1* plasmids fully rescued the AmB sensitivity of *hoi1Δ* mutants, whereas the binding-defective Hoi1(15A)-ENVY mutant showed no rescue. This suggests that the AmB sensitivity of *hoi1*Δ mutants is due to loss of Fmp27 and Hob2 recruitment to ER-PM contact sites.

**Figure 6:**
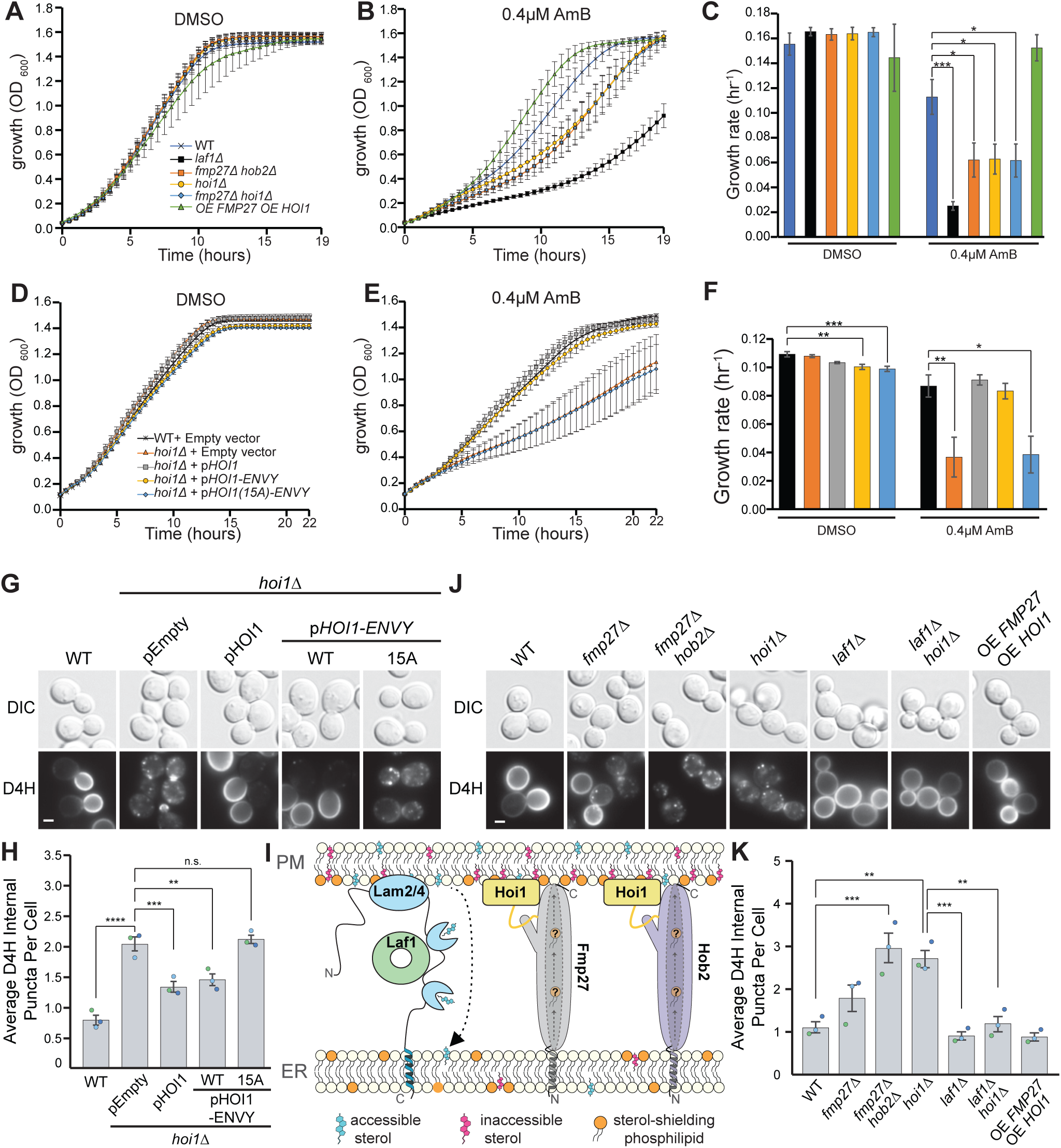
Fmp27 and Hoi1 have a shared role in lipid homeostasis. **(A-C)** *fmp27*Δ *hob2*Δ and *hoi1*Δ mutants are sensitive to amphotericin B (AmB). The growth of the indicated strains was monitored in the (A) absence or (B) presence of AmB by measuring the optical density (OD_600_) of cultures in 96-well plates. **(C)** Growth rate of strains from A, B. One-way ANOVA with Dunnett’s correction compared the growth rate of all strains to wild type (WT); n=3, p*<0.05, p***<0.001. Error bars indicate SEM. Only significant differences are indicated. OE, overexpressed. **(D-F)** Loss of the Hoi1-Fmp27 interaction confers sensitivity to amphotericin B (AmB). Growth was monitored in the (D) absence or (E) presence of AmB by measuring the optical density (OD_600_) of cultures in 96 well plates. (F) Growth rate of strains from D, E. One-way ANOVA with Dunnett’s correction comparing the growth rate of all strains to wild type (WT); n=4, p*<0.05, p**<0.01, p***<0.001. Error bars indicate SEM. Only significant differences are indicated. **(G)** Accessible sterols are mislocalized in *hoi1*Δ mutants. PM localization of the sterol reporter, mCherry-D4H, is rescued with plasmid-expressed wild type or tagged Hoi1, but not by the Fmp27 binding-defective 15A mutant. Fluorescent images of yeast cells expressing *TEF1pr-mCherry-D4H* (D4H) from the *FF21* locus. Bar = 2µm. **(H)** Automated quantitation of the average number of internal mCherry-D4H puncta per cell for strains in G. Individual data points are overlaid and coloured by replicate. One-way ANOVA with Tukey’s multiple comparison test; n = 3, cells/strain/replicate ≥ 364, **** = p<0.0001, *** = p<0.001, ** = p<0.01, n.s. = not significant. Error bars indicate SEM. **(I)** Schematic of proposed function of Hoi1, Fmp27 and Hob2 in regulating ergosterol distribution. In this model Fmp27 and Hob2, once recruited to the PM by Hoi1, transport sterol-shielding glycerophospholipids from the ER to the PM, which renders a pool of ergosterol in the cytosolic PM leaflet inaccessible. Loss of *HOI1*, or *FMP27* and *HOB2*, thus reduce levels of sterol-shielding lipids at PM, increasing the levels of accessible ergosterol that can readily be transported to the ER by Lam2/4 and their regulator, Laf1. **(J)** Accessible sterols are mislocalized to internal puncta in *hoi1*Δ and *fmp27*Δ *hob2*Δ mutants. This intracellular accumulation of sterols is lost in *laf1*Δ *hoi1*Δ double mutants. Wide-field images of yeast cells expressing *TEF1pr-mCherry-D4H* from the *FF21* locus. Bar = 2µm. **(K)** Automated quantitation of the average number of internal mCherry-D4H puncta per cell for strains in J. Individual data points are overlaid and coloured by replicate. One-way ANOVA with Tukey’s multiple comparison test; n = 3, cells/strain/replicate ≥ 408, *** = p<0.001, ** = p<0.01. Error bars indicate SEM.

Since AmB sensitivity can indicate either increased levels of accessible ergosterol in the outer PM leaflet, or general membrane integrity defects (Kamiński, 2014; Parsons et al., 2006) we assessed the distribution of accessible ergosterol within the cell by expressing the fluorescent sterol probe mCherry-D4H (Maekawa and Fairn, 2015) from a chromosomal locus. This probe, which contains the modified sterol-binding domain 4 (D4) of the *C. perfringens* theta-toxin, had a polarized distribution at the PM of WT yeast (Fig. 6G), with enhanced signal at the bud. However, mCherry-D4H was greatly reduced at the PM in the *hoi1Δ* mutant and instead accumulated in intracellular puncta (Fig. 6G-H). This phenotype was rescued by expressing WT Hoi1, but not a Hoi1(15A) mutant that is unable to interact with Fmp27. Mutations in the Hoi1 NT-C2 domain that disrupted its Fmp27-dependent PM targeting also resulted in D4H mislocalization (Fig. S3E,F). These results indicate that failure to recruit Fmp27 and Hob2 to ER-PM contacts leads to a redistribution of accessible ergosterol from the PM’s cytosolic leaflet to both the outer leaflet and internal compartments.

BLTPs are presumed to transport glycerophospholipids, rather than sterols. We hypothesized that loss of Fmp27 could indirectly affect sterol distribution by reducing the PM levels of sterol-shielding phospholipids (Maekawa and Fairn, 2015). This is predicted to increase sterol accessibility at the inner leaflet, allowing sterols to either flip to the outer leaflet or be recognized by the LAM family of lipid transporters for retrograde transport to the ER (Gatta et al., 2015). At the ER, sterols are diluted or esterified such that they are no longer recognized by the D4H probe, or transported to other organelles such as endosomes (Marek et al., 2020). This model, shown in Figure 6I, could explain why Fmp27 shows a strong functional correlation with the LAM2/4 sterol transporters and their regulator Laf1, and further explain why loss of Hoi1 causes accessible sterols to accumulate within the cell.

To test this model, we assessed the localization of mCherry-D4H in double mutant strains (Fig. 6J-K; Fig. S3G). As with *hoi1*Δ, the double *fmp27*Δ *hob2*Δ mutant accumulated D4H in intracellular puncta. In contrast, in a *laf1*Δ mutant, which is defective in PM-to-ER ergosterol transport, we observed an increase in mCherry-D4H intensity at the PM (Fig. S3G) and a loss of polarization between mother and daughter cells (Fig. 6J). Critically, combining *hoi1*Δ and *laf1*Δ mutations prevented the intracellular accumulation of mCherry-D4H and restored its PM localization. This rescue supports the model that loss of Fmp27 and Hob2 does not block sterol synthesis or initial delivery to the PM but instead liberates an accessible pool of ergosterol that is then cleared from the PM by Lam/Laf1-mediated retrograde transport.

### Fmp27 and Hoi1 orthologs in metazoans have shared phenotypes

Fmp27 and Hoi1 have homologs in metazoans that are not well characterized. AlphaFold2 predictions suggest that the nematode, fly and human orthologs form complexes that are structurally similar to Fmp27-Hoi1 (Fig. 7A-C). In each species, both the NT-C2 and tail regions of the Hoi1 homolog contact the BLTP2-like protein, with the tail interface limited to the BLTP2 “handle”. The functional importance of these proteins is highlighted in *C. elegans,* where mutations in the *HOI1* ortholog *sym-3* or the *LAF1* ortholog *sym-4* enhance the <u>P</u>harynx <u>in</u>gressed (Pin) phenotype of mutations in the apical extracellular matrix protein fibrillin (*fbn-1*) or the FBN-1 splicing factor *mec-*8 (Yochem et al., 2004; Kelley et al., 2015; Balasubramaniam et al., 2023). This synthetic interaction suggests these proteins help maintain the apical extracellular matrix needed to resist mechanical forces during pharyngeal development.

**Figure 7:**
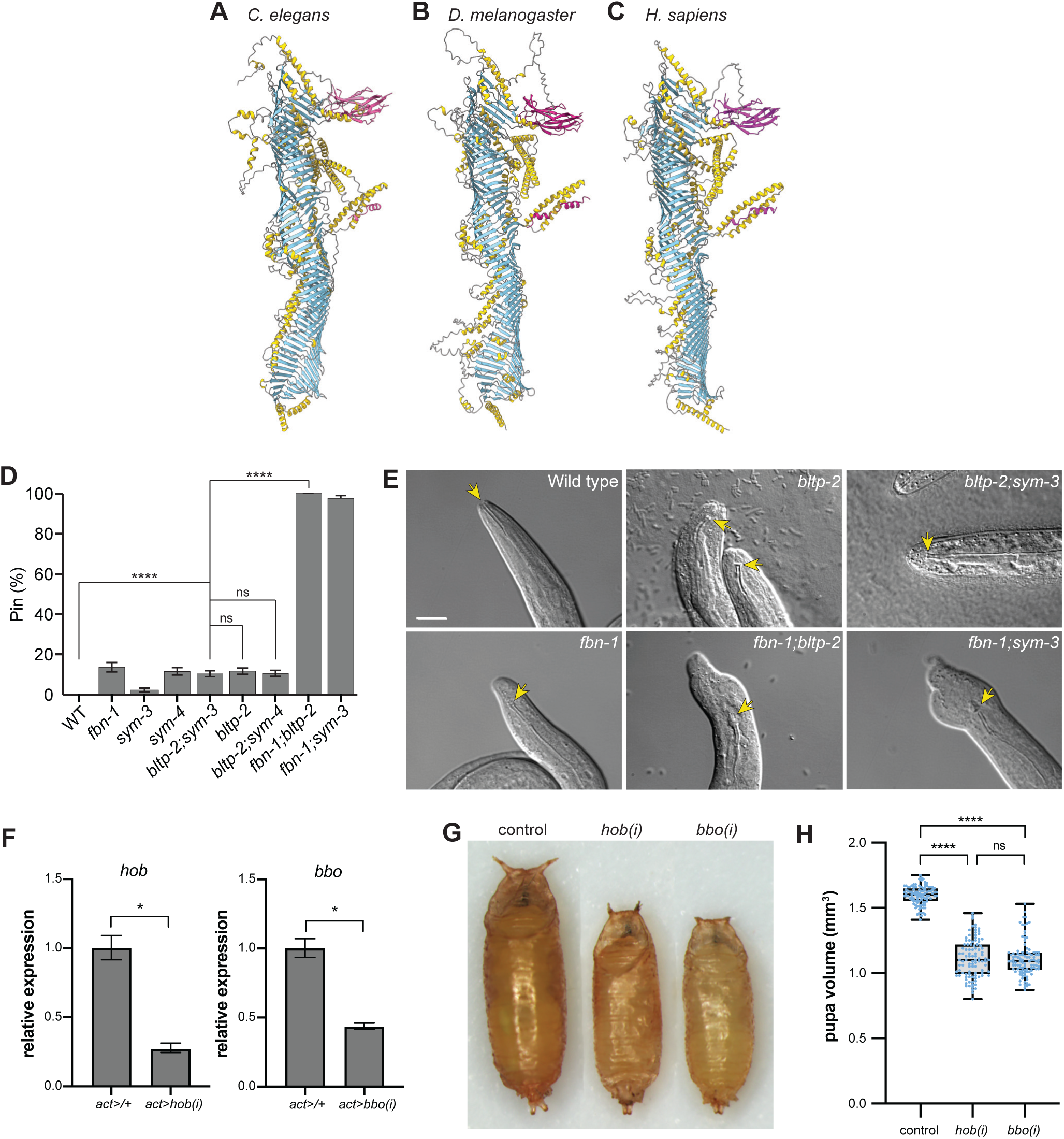
Higher eukaryote Fmp27 and Hoi1 orthologs have shared phenotypes. **(A-C)** ColabFold-predicted interactions between the Fmp27 and Hoi1 homologs from (A) *C. elegans* (B) *D. melanogaster* and (C) *H. sapiens*. Regions from Hoi1 homologs with pLDDT scores >70 (magenta) are shown to highlight confident interacting regions, whereas Fmp27 homologs are shown in their entirety. **(D)** Quantitation of the pharyngeal ingression (Pin) phenotype in embryo or L1 stage worms 24 hours post egg lay. Chi-square with Yates’ correction; **p<0.01, ****p<0.0001, ns = not significant. Error bars indicate standard error of proportion (SEP). **(E)** Representative images of Pin phenotype in L1 stage wild type, *fbn-1(tm290), bltp-2(ve727)*, *fbn-1;bltp-2, sym-3 and fbn-1;sym-3* mutants. Note the exaggerated head deformations in the *fbn-1;bltp-2* and *fbn-1;sym-3* double mutants. Yellow arrows mark the anterior end of the buccal cavity of the pharynx, which is abnormally ingressed in *bltp-2*, *fbn-1*, *bltp-2;sym-3*, *fbn-1;bltp-2*, *fbn-1;sym-3* mutant animals. **(F)** Validation of RNAi knockdown efficiency of *hob* and *bbo*. qPCR measurement of *hob* (left panel) and *bbo* (right panel) mRNA transcript levels in control and ubiquitous *hob-RNAi* or *bbo-RNAi* knockdown in whole animals at the onset of metamorphosis (puparium formation, 0 h PF) using the *act-GAL4* driver confirms that the RNAi constructs significantly reduce *hob* and *bbo* expression levels, respectively. We previously validated this *hobbit-RNAi* line using *tub-GAL4*. Relative expression, error bars, and statistics calculated by REST analysis. Independently isolated, biological triplicate samples measured for each genotype. *p<0.05. **(G)** Representative image of control (*act>/+*), ubiquitous *hob* knockdown (*act>hob(i)*), and ubiquitous *bbo* knockdown (*act>bbo(i)*) pupae. All three pupae were initially captured in the same image; animals were individually rotated and aligned post-acquisition to improve image aesthetics. **(H)** Body size quantified via pupal volume shows that size is significantly reduced upon ubiquitous knockdown of *hob* or *bbo*. Boxes outline the 25th to 75th percentiles; middle line indicates the median. Whiskers extend to the minimum and maximum values. Individual data points are overlaid; statistics calculated using ordinary one-way ANOVA. *n*=100 animals. ****p<0.0001, ns=not significant.

Based on our findings in yeast, we predicted that the uncharacterized *C. elegans bltp-2* protein works in the same pathway. Indeed, the *bltp-2* single mutant exhibited low levels of Pin similar to *fbn-1* single mutants, and *fbn-1;bltp-2* double mutants, like *fbn-1;sym-3* mutants, showed a dramatic enhancement of Pin and head malformation phenotypes (Fig. 7D,E). Importantly, the *bltp-2;sym-3* double mutant did not exhibit enhanced Pin. These results support a shared function for Fmp27 and Hoi1 orthologs in worms and further suggests a shared pathway with Laf1 that is conserved beyond yeast.

In *D. melanogaster*, the BLTP2 ortholog *hobbit* regulates body size and development through cell-autonomous control of insulin secretion (Neuman & Bashirullah 2018). The fly *HOI1* ortholog, *CG8671*, has not been extensively studied. Strikingly, we found that ubiquitous RNAi knockdown of *CG8671* phenocopied the small body size of ubiquitous *hobbit* knockdown (Fig. 7F-H). Furthermore, all *hobbit-RNAi* and *CG8671-RNAi* expressing animals arrested development during metamorphosis (n=100 per genotype). Due to the small body size and shared phenotypes with *hobbit*, we named the uncharacterized *CG8671* gene *bilbobaggins* (*bbo*) in honor of the small-statured character from J.R.R. Tolkien’s novels.

The striking phenotypic similarity between *hobbit* and *bbo* mutants suggests their protein products function together, like their orthologs in yeast. The small pupa size of *bbo* and *hobbit* mutants may result from secretory defects that are secondary to disruptions in lipid transport at membrane contact sites. Although we have not ruled out additional roles for these proteins at other internal membranes, our work suggests that the observed secretion defects may be the result of changes in lipid homeostasis at the PM, as we observe in yeast, which disrupt PM protein targeting and regulated exocytosis events.

## Discussion

We have found that the conserved BLTP2-like lipid transfer proteins Fmp27 and Hob2 are targeted to PM contacts through interactions with the previously uncharacterized protein Hoi1. Hoi1 forms an extensive interface with helical regions that extend from the central Fmp27 channel. The integrity of this interface is required for maintaining sterol homeostasis at the PM, suggesting that Hoi1-mediated targeting is important for the lipid transport function of Fmp27 and Hob2 at ER-PM contact sites.

### Hoi1 is the first identified adaptor for Fmp27

A variety of membrane adaptors mediate the targeting of Vps13 and Atg2, but until now no adaptor had been identified for BLTP2-like lipid transporters. Organelle-specific adaptors allow individual lipid transporters to target several different organelles in response to changing cellular conditions, and by reversibly detaching from membranes could regulate lipid flow. Although Fmp27 was also reported to be at ER-mitochondria contacts (Toulmay et al., 2022), the co-overexpression of both Hoi1 and Fmp27 led to their dramatic targeting to ER-PM sites. This suggests that Hoi1 is specific for the PM, and other adaptors may exist that target Fmp27 to other organelles.

BLTP2-like proteins lack recognizable domain insertions, like the VAB domain, that mediate interactions with organelle-specific adaptors in VPS13 proteins. We found that a prominent alpha-helical extension in Fmp27 (the “handle”), together with nearby helix and loop regions, interacts with helices and flanking sequences in the extended C-terminus of Hoi1. The paralogous Hob2 protein has a conserved arrangement of α-helical extensions, and similarly interacts with Hoi1. As ⍺-helical extensions decorate the hydrophobic channels of other members of the RBG superfamily (Neuman et al., 2022b; Levine, 2022), these may harbor binding sites for adaptors or regulators in other BTLPs.

How does Hoi1 target the PM? The Hoi1 N-terminal C2 domain, which is predicted to bind anionic membrane lipids in a calcium-independent manner (Zhang and Aravind, 2010), is necessary for the PM targeting of both Hoi1 and Fmp27, but does not associate strongly with membranes or bind Fmp27 on its own. The Fmp27 C-terminus may contribute low affinity interactions with membrane lipids that enhances targeting. In fact, the last 82 residues of the *Drosophila* Fmp27 homolog, Hobbit, are necessary for PM localization (Neuman et al., 2022a). The formation of a Fmp27-Hoi1 complex could thus create an extended platform for binding to PM lipids, which would explain why these proteins are mutually required for their PM targeting.

The Hoi1 NT-C2 domain could also bind to PM-localized proteins such as scramblases, to promote the redistribution of newly delivered lipids to the extracellular leaflet, which could prevent the buildup of lipids at their site of delivery and enable the coordinated expansion of both leaflets. In a similar manner, the binding of the structurally related Atg2 lipid transporter to its preautophagosomal adaptor, Atg18, relies on additional interactions with the lipid scramblase Atg9 (Gómez-Sánchez et al., 2018; Chowdhury et al., 2018). It is a tempting to speculate that the Hoi1 C-terminus is primarily responsible for initial targeting, whereas binding of the Hoi1 NTC2 domain promotes an Fmp27 orientation or interaction at the PM that facilitates lipid delivery. The BLTP2 and Hoi1 orthologs in humans, flies and nematodes are predicted to interact via these same interfaces, suggesting the complex and targeting mechanism is likely to be conserved. Understanding how these interactions regulate lipid flow across bridge-like transporters remains an important goal for future studies.

### Role of Fmp27, Hob2 and Hoi1 in ergosterol homeostasis

We determined that the loss of both yeast Hob-like lipid transporters, or their PM recruitment factor Hoi1, causes accessible sterol to be redistributed from the PM to intracellular compartments. This does not reflect a direct loss of ER-to-PM sterol transport. Instead, our data suggest that the loss of Fmp27 and Hob2 at ER-PM contact sites increases the pool of accessible sterol at the PM, which is then transported to intracellular compartments by members of the Lam family of retrograde sterol transporters (Gatta et al., 2015; Topolska et al., 2020).

If Fmp27 and Hob2 do not directly transfer sterols, what lipids do they transport at ER-PM contacts? Their similarity with other BLTP family members suggests they act on related lipid species. BLTPs are thought to preferentially bind glycerophospholipids (Kumar et al., 2018; Valverde et al., 2019; Kang et al., 2024 *Preprint*), and can transport PE, PC and PS between liposomes (Kumar et al., 2018; Osawa et al., 2019; Valverde et al., 2019; Hanna et al., 2022). Recent work suggests that yeast and nematode BLTP1-like proteins transport unsaturated phospholipids to the PM (John Peter et al., 2022; Wang et al., 2022), and may also transport phosphatidylethanolamine (PE) to the ER (Toulmay et al., 2022). This suggests Fmp27 and Hob2 also transport glycerophospholipids, and that loss of this transport indirectly alters ergosterol accessibility at the PM.

The accessibility of ergosterol is modulated by its co-distribution with other lipids that can shield it from proteins such as lipid transporters, biosynthetic enzymes, or synthetic probes (Maekawa and Fairn, 2015; Kishimoto et al., 2021). In yeast, ergosterol distribution at the PM is asymmetric, with roughly 80% in the cytosolic leaflet and 20% in the outer leaflet (Solanko et al., 2018). Altering the transbilayer distribution of sterol-shielding glycerophospholipids causes a reduction in total sterol levels at the PM and a relative increase in the proportion of sterols in outer versus cytosolic PM leaflets (Solanko et al., 2018; Kishimoto et al., 2021; Maekawa and Fairn, 2015). The similar redistribution of ergosterol we observe in *fmp27 hob2* and *hoi1* mutants is consistent with the model that Fmp27 and Hob2 maintain sterol organization by delivering specific glycerophospholipids to the PM.

Alterations in levels of accessible sterol in yeast could also result from homeostatic cellular mechanisms that maintain membrane fluidity. Organisms respond to temperature changes by compensatory alterations in cholesterol levels, PE:PC ratios, and acyl chain length or saturation (Renne and de Kroon, 2018; Wu et al., 2023). Indeed, a recent study shows that loss of BLTP2 proteins in yeast, worms or human cells causes increased membrane fluidity and reduced PM levels of PE specifically in response to cold (Banerjee et al., 2024 *Preprint*). Although sterols are not an abundant class of membrane lipids in flies or worms, which are cholesterol auxotrophs, both human and insect cells respond to changes in sterol levels by modulating levels of PE (Rauthan and Pilon, 2011; Dawaliby, 2016). Determining if the observed changes in a specific lipid are due to defects in the transport of that lipid, or are indirect consequences of lipid homeostasis mechanisms, will require additional studies under a variety of conditions that alter membrane composition or fluidity.

### Conservation of the Fmp27-Hoi1 complex in higher eukaryotes

Our work in flies and nematodes suggests the Fmp27-Hoi1 complex is conserved in metazoans and provides evidence for a shared function in secretion. We previously determined that loss of the *Drosophila FMP27* homolog, *hobbit*, results in cell-autonomous defects in regulated exocytosis, which was the first described function of an animal BLTP2 ortholog (Neuman and Bashirullah, 2018; Neuman et al., 2022a). Knockdown of the *HOI1* ortholog, *bbo*, phenocopies loss of *hobbit*, suggesting that both proteins are functionally linked. In *C. elegans,* reduced levels of this complex enhances a developmental defect that may be linked to the reduced secretion of different apical extracellular matrix proteins (Kelley et al., 2015; Balasubramaniam et al., 2023). Are secretion defects in these mutants related to the PM sterol homeostasis defects seen in yeast? Lipids such as cholesterol, phosphoinositides and glycerophospholipids are important for vesicle priming and fusion at the PM (Zhang et al., 2009; Chasserot-Golaz et al., 2010; Ammar et al., 2013; Tanguy et al., 2016; Yang et al., 2021), suggesting lipid imbalances at the PM could underlie the observed intracellular trafficking and secretion defects of *hobbit* and *bbo* mutants.

Mutations in the *C. elegans* homologs of Fmp27, Hoi1 and Laf1 cause strikingly similar phenotypes, suggesting all three proteins work in a shared pathway (Yochem et al., 2004; Kelley et al., 2015; Balasubramaniam et al., 2023). The recent demonstration that the human BLTP2 protein colocalizes with the Laf1 homolog WDR44 at a tubular endosomal network in HeLa cells, and that both proteins negatively regulate ciliogenesis (Parolek and Burd, 2024), further highlights the functional connections between these proteins. The role of the two human Hoi1 homologs, EEIG1 and EEIG2 (Choi et al., 2013; Jeong et al., 2020), in BLTP2 targeting and function remains to be explored. BLTP2 overexpression is associated with reduced survival of cancer patients (Song et al., 2006; Liu et al., 2014; Zhong et al., 2018) and a recent study finds that missense mutations in BLTP2 are linked to neurodevelopmental disorders (Ng et al., 2024 *Preprint*). Further studies will be required to better understand how these proteins regulate lipid transport by BLTP2.

## Materials and Methods

### Yeast Strains and Plasmids

Yeast strains and plasmids, and primers used in the generation of PCR products for strain and plasmid construction, are listed in Supplemental Table 2. Plasmids were made by homologous recombination in yeast by co-transforming linearized plasmids with PCR products, recovering in *Escherichia coli* and sequencing. Yeast strains were made by homologous recombination-based integration of PCR products (Janke et al., 2004; Longtine et al., 1998) unless otherwise stated. Gene deletions and promoter exchanges were confirmed by PCR, and epitope tags were confirmed by microscopy, western blot, and/or PCR. The Markerless Yeast Localization and Overexpression (MYLO) CRISPR-Cas9 toolkit (Bean et al., 2022) was used to generate strains with stable integration of RPL18B-ymNeonGreen-2xPH(PLCδ) or TEF1-mCherry-D4H. Briefly, yeast were co-transformed with two linearized plasmids: the first providing donor DNA for repair and the second providing Cas9 and gRNA targeting the designated genomic locus. Genomic integration events that restored the selectable marker by repair of the Cas9-gRNA plasmid by homologous recombination at a split-selection marker were confirmed by microscopy. CRISPR-Cas9-based genome-editing approaches were also used to generate *fmp27* point mutants and introduce internal ymNeonGreen-tags. Custom linearized Cas9-gRNA plasmids targeting *FMP27* sequences were co-transformed with PCR-amplified donor DNA. Validated CRISPR-Cas9 edited strains were validated by sequencing and grown without selection to lose gRNA plasmids and restore auxotrophy.

### Microscopy Image Acquisition and Processing

Yeast cells were cultured overnight in synthetic minimal media (SD) (0.68% [w/v] yeast nitrogen base, 2% [w/v] ammonium sulfate, amino acid supplements) at 30 °C, diluted and grown to logarithmic phase. Cells were transferred to 96-well glass-bottom plates (Eppendorf or Cellvis) pre-treated with concanavalin A (Sigma-Aldrich). After 4 minutes, wells were aspirated to remove non-adherent cells and fresh SD media was added for imaging. Widefield images were acquired at room temperature on a Leica TCS SP8 Microscope (Leica Microsystems, Wetzlar, Germany) with a high-contrast Plan Apochromat 63×/1.30 Glyc CORR CS objective (Leica Microsystems, Wetzlar, Germany) and an ORCA-Flash4.0 digital camera (Hamamatsu Photonics, Shizuoka, Japan). Fluorescent and DIC images were acquired and processed with MetaMorph 7.8 software (MDS Analytical Technologies, Sunnyvale, California), with linear intensity scale changes uniformly applied to all images of a given fluorophore. For very bright or very dim signals that could not be identically scaled, custom settings were applied to each image to show protein localization. Confocal images were acquired with Leica Application Suite X (LASX) 3.5.7 software (Leica Microsystems, Wetzlar, Germany) (490nm for NeonGreen/ENVY/EGFP/sfGFP and 570nm for mScarletI) and deconvolved using Huygens Essentials software (Scientific Volume Imaging, Hilversum, Netherlands). Representative confocal images were individually adjusted for brightness and contrast in Photoshop CC 2022 (Adobe, Mountain View, California).

### Quantitation of Imaging Data

Cortical recruitment of overexpressed Fmp27-Envy by RPH-Hoi1 chimeras was quantified by blind counting of at least 100 cells in three independent replicates using the Fiji Cell Counter tool (Schindelin et al., 2012). For representative line scans to visualize colocalization, profiles of the cell cortex were extracted and linearized using Fiji. Fluorescent signal intensities were normalized to absolute values and plotted. To determine PM to nER intensity ratios, cells with clearly separated nuclear and cortical ER were identified using the ER marker Sec63-mScarletI. Selections were made around the cell cortex (PM) and the nER in Sec63-mScarletI images and propagated to the corresponding Fmp27^NeonGreen or Hoi1-ENVY images using Fiji. Fluorescent signal intensities across each pixel were extracted and the average intensity used to compute the PM to nER intensity ratio for each cell. To determine PM to cytosol intensity ratios, selections were made around the cell cortex (PM) or within the cell in the DIC channel and propagated to Hoi1-mScarletI images. Fluorescent signal intensities across each pixel were extracted and the average intensity used to compute the PM to cytosol intensity ratio for each cell. Statistical analyses were carried out using GraphPad Prism 9.1.0 (GraphPad Software; San Diego, California). Data distribution was assumed to be normal, but not formally tested.

For automated quantitation of the number of mCherry-D4H internal puncta per cell and mCherry-D4H area at the PM, the mCherry-D4H accessible sterol reporter and the NeonGreen-2xPH(PLCδ) PM reporter were integrated at separate loci, and fluorescent images were captured with GFP, RFP and BFP filter sets. Images were analyzed using a custom MetaMorph 7.8 journal. For all strains analyzed, live cells were identified in the BFP channel using the Count Nuclei function. Dead cells and extracellular noise in GFP, RFP and BFP channels were identified with iterative Count Nuclei functions using high intensity thresholds and masked with the Arithmetic function. A final Count Nuclei computation was used on the masked images to generate a live cells mask. The PM was identified with the Morphological Top Hat function in the GFP channel using the NeonGreen-2xPH(PLCδ) marker, selected with the Count Nuclei function, inverted, and eroded. A PM-masked image was generated by combining the live cells mask and eroded PM image using the Arithmetic function. An inversed masked image of only the PM was similarly generated by combining the live cells mask with an inverted eroded PM mask using the Arithmetic function. mCherry-D4H signals were identified with the Granularity function and combined with the live cells mask to identify mCherry-D4H signals in live cells. Using the PM masked image, an additional Granularity computation was used to identify intracellular mCherry-D4H puncta. The number of intracellular puncta per cell was computed by dividing the number of intracellular puncta within an image by the number of live cells identified in that image. mCherry-D4H signals at the PM were identified with the Granularity function on the PM-only masked image and the cumulative size of all identified regions within an image computed and divided by the number of live cells identified in that image. Statistical analyses were carried out GraphPad Prism 9.1.0. Data distribution was assumed to be normal, but not formally tested.

### Western blot and co-immunoprecipitation

For western analysis of protein stability, yeast cells were cultured overnight in complete yeast peptone dextrose media (YPD) (1% [w/v] yeast extract, 2% [w/v] Bacto-Peptone, 2% [w/v] glucose, 0.032% [w/v] tryptophan) at 30 °C, then diluted and grown to logarithmic phase. 10 OD_600_/mL equivalent of cells were harvested, washed, and stored at −80 °C for 20 minutes. Cells were resuspended in 100μL Thorner buffer (40mM Tris-Cl pH 7.0, 8M urea, 5% SDS, 1% β-mercaptoethanol and 0.4 mg/mL bromophenol blue) and lysed by vortexing with 0.5 mm diameter glass beads three times at 45 second intervals with 1 minute incubation on ice between vortexes. Proteins were denatured at 70 °C for 5 minutes. For Fmp27-ENVY, 1 OD_600_/mL equivalent of cell lysate was separated on 7% SDS-PAGE gels. For Hoi1-ENVY and Pgk1, 0.4 and 0.5 OD600/mL equivalent of cell lysate respectively was separated on 10% SDS-PAGE gels.

For co-immunoprecipitation experiments, 70 OD_600_/mL equivalent of cells were harvested, washed, and stored at −80 °C for 20 minutes before vortexing in 500μl lysis buffer (50 mM HEPES pH 7.4, 1% Tween-20, 150 mM NaCl, 1 mM EDTA, 1 mM PMSF, and 1X yeast/fungal Protease Arrest) with 0.5 mm diameter glass beads. 50μl of lysate was set aside and resuspended in 4X Laemmli Sample Buffer (240 mM Tris-Cl pH 6.8, 8% SDS, 40% glycerol, 0.004% bromophenol blue) with 8M urea and 10% β-mercaptoethanol. The remainder of each lysate was incubated with polyclonal rabbit anti-GFP (EU2; Eusera) for 1 h followed by Protein A-Sepharose beads (GE Healthcare) for another hour at 4°C. Washed beads were resuspended in 50μl of Thorner buffer and proteins eluted by heating at 100°C for 5 min.

After separation on SDS-PAGE gels, proteins were transferred to nitrocellulose membranes (50-206-3328; Fisher Scientific) and blotted with either mouse monoclonal α-GFP (11814460001; Roche), HA.11 (MMS-101R; Covance) or α-PGK1 (22C5D8; Thermo Fisher Scientific) antibodies, followed by horseradish peroxidase-conjugated polyclonal goat α-mouse (115–035-146; Jackson ImmunoResearch Laboratories) antibodies. Blots were developed with ECL chemiluminescent reagents (RPN2106, Cytiva) and exposed to Amersham Hyperfilm ECL (CA95017-653L; VWR). Densitometry of scanned films was performed in Fiji. Statistical analysis was carried out using GraphPad Prism 9.1.0. Data distribution was assumed to be normal, but not formally tested.

### Structural Modelling and Analysis

Prediction of protein-protein interactions was performed by the ColabFold AlphaFold2-Multimer MMseq2-driven homology search software (Mirdita et al., 2022; https://colab.research.google.com/github/sokrypton/ColabFold/blob/main/AlphaFold2.ipynb) with default settings. The pLDDT threshold of 70 was applied using PyMOL (The PyMOL Molecular Graphics System, Version 3.0 Schrödinger, LLC), and the analysis of predicted contact residues was performed using the ‘Protein interfaces, surfaces and assemblies’ service PISA at the European Bioinformatics Institute. (http://www.ebi.ac.uk/pdbe/prot_int/pistart.html) (Krissinel and Henrick, 2007) on the thresholded model. Conservation analyses were performed on the ConSurf web server (consurf.tau.ac.il; Ashkenazy et al., 2106). Figures were generated in UCSF Chimera version 1.16.0 or UCSF ChimeraX version 1.8 (Pettersen et al., 2004; Pettersen et al., 2021 University of California San Francisco Parnassus Campus) and assembled in Adobe Illustrator 2022 (Adobe, Mountain View, California).

### Chemicogenomic Dataset and Gene Ontology Analysis

Genome-wide Pearsons correlation scores and associated p values were computed in RStudio (RStudio Team, Boston, Massachusetts) using the homozygous deletion profiling (HOP) chemicogenomics dataset (Hoepfner et al., 2014). The network of genes showing a genetic correlation with FMP27 with a p-value <0.01 was generated in Cytoscape version 3.5.1 (Shannon et al., 2003). The correlation values were plotted in Microsoft Excel 2022 (Microsoft, Redmond, Washington) and formatted in Adobe Illustrator 2022 (Adobe, Mountain View, California). Gene Ontology (GO) functional enrichment analysis was performed using the Gene Ontology Consortium (Ashburner et al., 2000; The Gene Ontology Consortium et al., 2021)

Enrichment Analysis tool (Mi et al., 2019). The *Saccharomyces cerevisiae* reference was used and GO biological process complete annotation data set selected for analysis, using Fisher’s Exact test and Bonferroni correction for multiple testing.

### Growth assays

For plate-based drug sensitivity assays, overnight cultures were diluted and grown 4 hours to a concentration of 1-2 OD_600_/mL. 10-fold serial dilutions were spotted onto YPD plates or YPD supplemented with amphotericin B (Sigma). For liquid culture growth assays, the method from Hung et al., 2018 was adapted. Briefly, 100µL of 0.2 OD_600_/mL log phase cultures in either YPD or synthetic dextrose (SD) media were transferred to a 96 well plate. Amphotericin B, diluted in the same media to twice the desired final concentration, was then added to each well, and mixed by pipetting. OD_600_ readings were taken every 30 minutes, for at least 19 hours, with intermittent orbital shaking at 360 rpm for 2 minutes. To minimize evaporation, the plate was covered with a lid, which was treated with 0.05% Triton-X and 20% ethanol and air dried to prevent condensation and removed for each reading. Blank OD_600_ readings were subtracted from the sample OD_600_ readings, technical replicates were averaged and OD_600_ vs time (in hours) was plotted. OD_600_ values corresponding to the log phase of growth from three independent experiments were used to calculate average growth rate. Statistical analysis was carried out using GraphPad Prism 9.1.0. Data distribution was assumed to be normal, but not formally tested.

### C. elegans strains, maintenance and quantification of Pin phenotype

*C. elegans* strains were cultured using standard techniques on nematode growth media-lite (NGM-l) plates, with *E. coli* OP50 as the food source (Brenner, 1974). All experiments were carried out at 20°C unless otherwise specified. Worm strains used in this study included the following: N2, WY1034 [*fbn-1(tm290) III; fdEx249 (sur-5::GFP; fbn-1(+)-fosmid wrm0635cH08)*], SP2231 [*sym-3(mn618) X*], SP2232 [*sym-4(mn619) X*], RG3227 [*bltp-2(ve727[LoxP + myo-2p::GFP::unc-54 3’UTR + rps-27p::neoR::unc-54 3’UTR + LoxP]) I*], STE143 [*bltp-2(ve727); sym-3(mn618); mnEx169 (sym-3(+);sur-5::GFP)*], STE144 [*bltp-2(ve727); sym-4(mn619); fdEx226 (sym-4(+); sur-5:GFP; Phsp-16::peel-1)*], STE145 [*bltp-2(ve727); fbn-1(tm290); fdEx249*], and WY1058 [*fbn-1(tm290) III; oyIs44 V; sym-3(mn618) X; fdEx249*]. STE143, STE144, STE145 were generated by standard genetic crossing of respective single mutants with balanced strains WY1034, WY893 [*mec-8(u74) I; sym-3(mn618) X; mnEx169 (sym-3(+);sur-5::GFP)]*, and *WY969 [mec-8(u74) I; sym-4(mn619) X; fdEx226 (sym-4(+); sur-5:GFP; Phsp-16::peel-1)*] (Kelley et al., 2015). RG3227 was generated by the CGC using CRISPR-Cas9 genome editing (Au et al., 2019).

To obtain embryos, synchronized adults were allowed to lay eggs for 8 hours and hatched overnight at room temperature. Offspring in the form of eggs and L1 were collected in M9+0.01% Tween and immobilized in 12.5 mM levamisole (Sigma L9756) on 2% (w/v) agarose pads. Percent Pin was scored in 3-fold or older embryos or L1-stage larvae under a Leica SP8X microscope at 100× magnification. Image acquisition was carried out with MetaMorph software (Molecular Devices). For each experiment, at least three independent trials were performed, totalling at least 120 offspring per genotype. For balanced strains (WY1034, STE143, STE144, STE145, WY1058), balanced GFP-bearing parents were used for egg-lay, and only non-GFP offspring were scored for the Pin phenotype.

### Drosophila husbandry, stocks, and body size analysis

The following stocks were obtained from the Bloomington *Drosophila* Stock Center: *w^1118^* (RRID:BDSC_3605), *act-GAL4* (RRID:BDSC_3954) and *CG8671-RNAi* (RRID:BDSC_41965). The *hobbit-RNAi* stock was previously described (Neuman & Bashirullah 2018). All experimental crosses were grown on standard cornmeal-molasses media in density-controlled bottles or vials kept in an incubator set to 25°C. Body size quantification was performed as previously described (Neuman and Bashirullah, 2018); briefly, the length and width of each pupa was measured using Adobe Photoshop CS6 and the resulting measurements were used to calculate pupa volume using a published formula (Delanoue et al. 2016). Pupa images were captured on an Olympus SXZ16 stereomicroscope coupled to an Olympus DP72 digital camera with DP2-BSW software. All pupae pictured together were originally captured in the same image but were individually rotated and aligned post-acquisition to improve image aesthetics.

### Quantitative real-time PCR (qPCR)

qPCR was performed as previously described (Ihry et al., 2012). Total RNA was isolated from whole animals at the onset of metamorphosis/puparium formation (0 h PF) using the Qiagen RNeasy Plus Mini Kit. 100 ng of total RNA was used to synthesize cDNA using the SuperScript III First-Strand Synthesis System (Invitrogen). Samples were collected and analyzed in biological triplicate. qPCR was performed on a Roche LightCycler 480 with LightCycler 480 SYBR Green I Master Mix (Roche). Relative Expression Software Tool (REST) was used to calculate relative expression and *p*-values; this software calculates standard error using confidence intervals centered on the median and determines *p*-values for relative expression ratios using integrated randomization and bootstrapping methods (Pfaffl et al., 2002). Expression was normalized to the reference gene *rp49*; primers for *rp49* were previously published (Ihry et al., 2012). The *hob* qPCR primers were previously published (Neuman and Bashirullah, 2018). New primers for *CG8671/bbo* were designed using FlyPrimerBank (Hu et al., 2013).

## Supplemental material

Figure S1 (related to Figure 3) provides additional figures examining the role of the Hoi1 N and C-terminal domains. Figure S2 (related to Figure 5) provides AlphaFold confidence measures. Figure S3 (related to Figure 6) shows additional measures of ergosterol distribution in mutants. Table S1 provides a PISA analysis of residues at the Fmp27-Hoi1 interface. Table S2 provides a list of strains, plasmids and primers used in this study.

## Acknowledgments

We thank Dr. Bjorn Bean, Dr. Malcom Whiteway and Dr. Vincent Martin (Concordia University, Montréal, Canada) for the MYLO CRISPR-Cas9 genome-editing toolkit. We thank Dr. Maya Schuldiner (Weizman Institute of Science, Rehovot, Israel) and Dr. Chris Stefan (University College London, London, United Kingdom) for providing lipid reporter plasmids, and thank Dr. Wanda Kukulski (Universitat Bern, Switzerland) for yeast strains. We additionally thank Omer Riyadh for providing a custom R script. We thank Dr. D. S. Fay for *C. elegans* strains and for helpful discussions.

## Author contributions

Funding acquisition, Project administration, Resources, Supervision: E.C, S.T, A.B. Investigation, Methodology, Visualization: S.K.D, V.S, H.E., E.C., M.D., S.D.N, J.Y. Conceptualization, Writing – original draft, Writing – review & editing: E.C, S.T, A.B., S.K.D, V.S, H.E., S.D.N, J.Y.

## Abbreviations

aa: Amino acid
GFP: Green fluorescent protein
HA: Hemagglutinin
mScI: mScarletI
OE: Over-expressed
PAE: Predicted aligned error
PM: plasma membrane
pLDDT: Predicted local distance difference test
RFP: Red fluorescent protein
WCL: whole cell lysate
WT: Wild-type

---

**Supplemental Figure 1:**
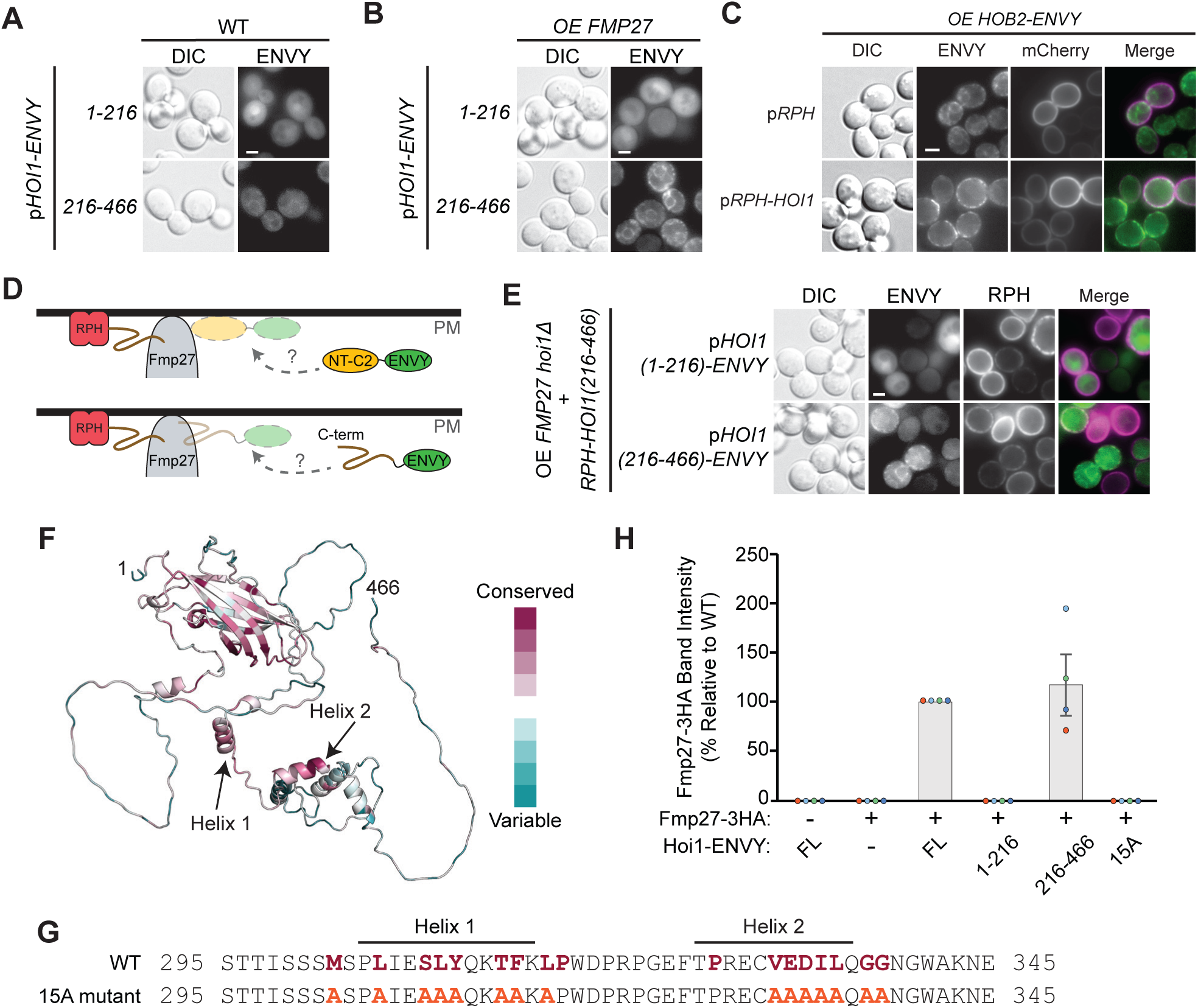
Fmp27 recruitment by the Hoi1 C-terminal domain. **(A, B)** The Hoi1 C-terminus is recruited to membranes in an *FMP27*-dependent manner. Hoi1-Envy N-terminal (1-216) or C-terminal (216-466) fragments were overexpressed in cells in the (A) absence or (B) presence of co-overexpressed *FMP27*. **(C)** Overexpressed (OE) Hob2-Envy is recruited to cortical ER-PM contacts by the RPH-Hoi1 chimera, but not by the RPH module alone. Representative widefield images are shown. Bar = 2µm. **(D, E)** Overexpressed Fmp27 that is artificially recruited to the PM by the RPH-Hoi1(216-466) chimera does not recruit the Hoi1(1-266)-Envy N-terminal domain. A schematic illustrating the experiment is shown in (D); widefield images of cells expressing the indicated proteins are shown in (E). Bar = 2µm. **(F)** AlphaFold predicted model of Hoi1 with residues colored by conservation scores. **(G)** Highly conserved residues within the region of the Hoi1 C-terminal tail that is critical for Fmp27 recruitment are shown in pink; the 15 residues that were changed to alanine in the 15A mutant are shown in orange. **(H)** The Hoi1(15A) mutant does not bind Fmp27. Quantitation of the co-immunoprecipitation experiment from Fig 3I, showing the levels of Fmp27-3HA that co-purify with ENVY-tagged wild type (WT) and mutant forms of Hoi1 in an anti-GFP pulldown. Individual data points are overlaid and coloured by replicate, n=4.

**Supplemental Figure 2:**
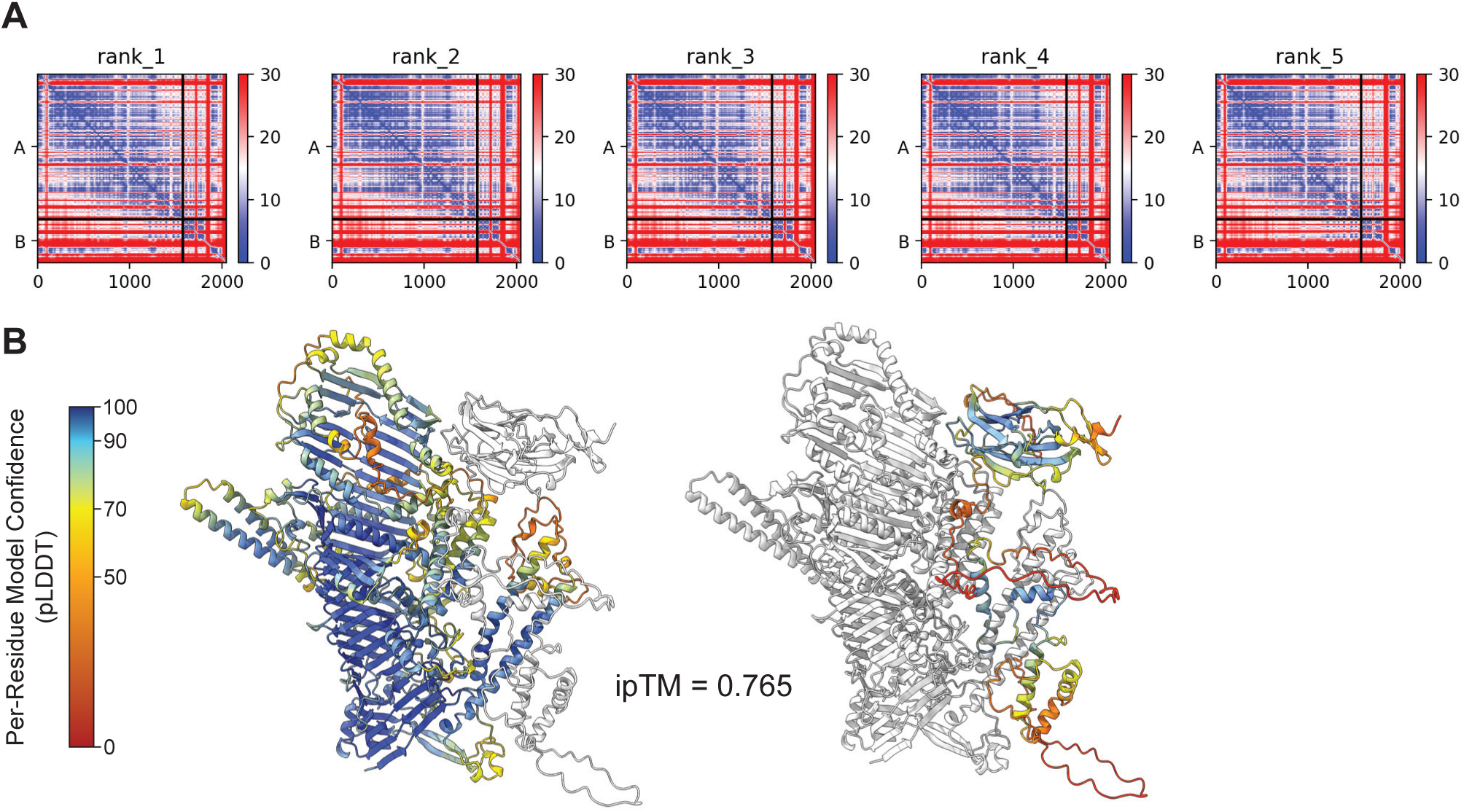
AlphaFold confidence measures. **(A)** ColabFold-generated predicted aligned error (PAE) plots for the top 5 ranked models of the Fmp27^1051-2628^ (Chain A) and Hoi1^1-466^ (Chain B) interaction. Low predicted aligned error indicates high confidence interactions. **(B**) The ColabFold-predicted interface between Fmp27 (1051-2628) and Hoi1, showing all residues for the top-ranked model. Left image shows Fmp27 colored in gray, with Hoi1 colored by per-residue pLDDT values. Right image shows Hoi1 colored in gray, with Fmp27 colored by per-residue pLDDT values. ipTM: interface predicted template modelling score.

**Supplemental Figure 3:**
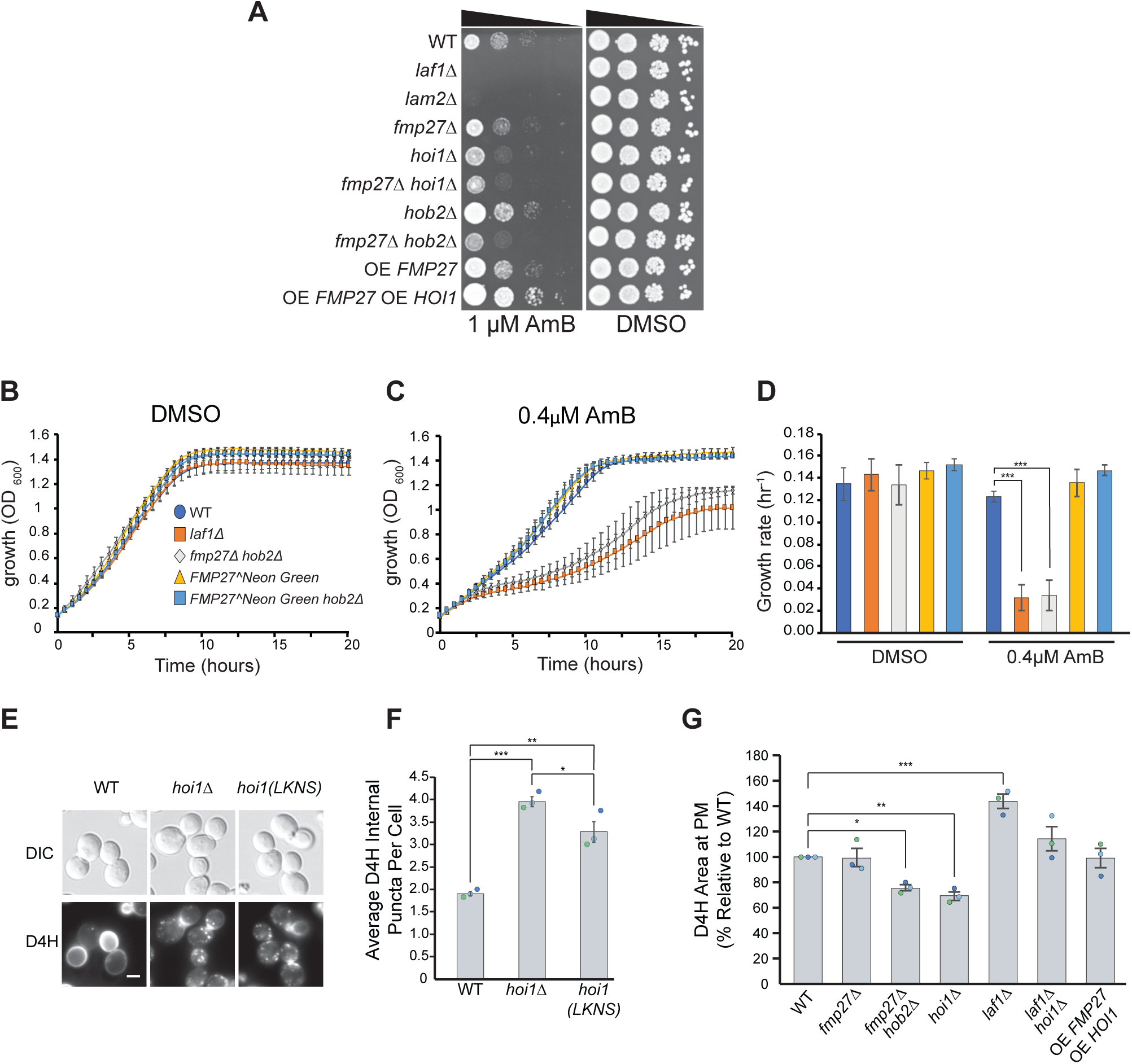
Additional measures of ergosterol distribution. **(A)** 10-fold serial dilutions of the indicated yeast strains grown for 3 days at 30°C on solid YPD media supplemented with DMSO or 1μM amphotericin B (AmB) in DMSO. **(B-D)** Fmp27 internally tagged with NeonGreen is fully functional. The growth of the indicated strains was monitored in the absence (B) or presence (C) of AmB by measuring the optical density (OD_600_) of liquid cultures in 96-well plates. (D) Growth rate of strains from B, C. One-way ANOVA with Dunnett’s correction comparing the growth rate of all strains to wild type (WT); n=3, p***<0.001. Error bars indicate SEM. Only significant differences are indicated. **(E, F)** Mutation of Hoi1 NT-C2 residues that contact Fmp27 cause mislocalization of the Cherry-D4H reporter. (E) Fluorescent images of yeast cells expressing *TEF1pr-mCherry-D4H* from the *FF21* locus. Bar = 2µm. (F) Quantitation of strains shown in E. One-way ANOVA with Tukey’s multiple comparisons test; n=3, cells/strain/replicate ≥450. Error bars indicate SEM. **(G)** Quantitation of the average area of mCherry-D4H at the PM per cell for strains in Figure 6J. Individual data points are overlaid and coloured by replicate. One-way ANOVA with Tukey’s multiple comparison test; n = 3; cells/strain/replicate ≥ 408; *** = p<0.001; ** = p<0.01; * = p<0.5. Error bars indicate SEM.

